# Oxidative Stress Susceptibility, Complement Dysregulation, and Metabolic Reprogramming in *CFH* Y402H Patient-Derived Choriocapillaris Endothelial Cells

**DOI:** 10.64898/2026.06.18.733111

**Authors:** Agata Rozanska, Jasenka Guduric-Fuchs, Varun Pathak, Pietro M. Bertelli, Matthew D. Teasdale, Harriet Denton, Diya Bhattacharyya, Rafiqul Hussain, Jonathan Coxhead, Joe Kelk, Kevin Marchbank, David Kavanagh, Marzena Kurzawa-Akanbi, Lyle Armstrong, Reinhold Medina, Majlinda Lako

**Affiliations:** Biosciences Institute, Newcastle University, UK; The Wellcome Wolfson Institute for Experimental Medicine, Queen’s University, Belfast; Translational and Clinical Research Institute, Newcastle University, UK; Department of Eye and Vision Sciences, Institute of Life Course and Medical Sciences, University of Liverpool; NIHR Newcastle Biomedical Research Centre (BRC), Newcastle upon Tyne, UK; BSU affiliation: Bioinformatics Support Unit, The Faculty of Medical Sciences, Newcastle University, Newcastle upon Tyne NE2 4HH, UK

**Keywords:** Age-related macular degeneration (AMD), complement factor H (CFH), choriocapillaris endothelial cells (CECs), oxidative stress, lipid metabolism

## Abstract

Age-related macular degeneration (AMD) is a leading cause of central vision loss. Immunofluorescence and gene expression studies in human donor eyes have shown that choriocapillaris endothelial cells (CECs) are lost before retinal pigment epithelium (RPE) degeneration, leaving extracellular matrix–filled empty lumens known as “ghost vessels.” To investigate disease mechanisms, we generated CECs from patient-specific induced pluripotent stem cells (iPSCs) carrying the high-risk *CFH* Y402H variant and CRISPR-Cas9–corrected isogenic controls. The iPSC-derived CECs expressed canonical endothelial markers, formed fenestrations, maintained barrier function, and assembled capillary-like structures. Although baseline metabolism was preserved, Y402H CECs showed heightened sensitivity to hydroquinone-induced oxidative stress, with increased cytotoxicity and deposition of the complement membrane attack complex (C5b–9). RNA sequencing revealed oxidative stress–driven upregulation of lipid biosynthesis, mTORC signalling, endothelial-to-mesenchymal transition, and angiogenic pathways, alongside an imbalance in the complement pathway. These findings demonstrate that the *CFH* Y402H polymorphism increases CEC vulnerability to environmental stress, linking complement dysregulation and metabolic reprogramming to choriocapillaris dysfunction in AMD and highlighting CECs as a potential therapeutic target.

## Introduction

Age-related macular degeneration (AMD) is a chronic condition that gradually degrades the retina’s central macular zone, eventually causing loss of central vision (Mitchell et al., 2018). Recognised by the World Health Organisation as the third most prevalent cause of vision impairment worldwide, its reach is significant: roughly 8.06 million people were affected in 2021. This number is expected to climb to 21.34 million by 2050, creating a significant global socioeconomic impact (GBD 2021 Global AMD Collaborators, 2025). The hallmark signs of AMD include the buildup of drusen, which are fatty deposits, alongside abnormalities within the retinal pigment epithelium (RPE). This disease typically advances through three primary phases, beginning with an often-asymptomatic early stage identified by the presence of medium-sized drusen measuring between 63 and 125 µm in the macula. It then transitions into an intermediate stage characterised by larger drusen exceeding 125 µm and noticeable changes in retinal pigmentation, eventually reaching the late stage where the condition manifests as either “dry” AMD, known as geographic atrophy, or “wet” AMD, referred to as neovascularisation (Mitchell et al., 2018). In the advanced stages of the disease, death of retinal photoreceptors, RPE and choroid endothelial cells (CECs) is common. Loss of endothelial cells from the choriocapillaris is one of the earliest detectable events in AMD (Mullins et al., 2011, Whitmore et al., 2013).

The choroid is a highly specialised vascular network, lying just beneath the RPE, between the Bruch’s membrane and the sclera. It is composed of three layers: the innermost layer (choriocapillaris), middle (Sattler’s), and outer layer (Haller’s layer). The choriocapillaris is a continuous layer of capillaries with surrounding pericytes organised in a lobular arrangement and is the principal blood supply to RPE and photoreceptors. Importantly, the choriocapillaris supplies the central retinal region responsible for high acuity vision (fovea), which lacks retinal blood vessels (Margolis and Spaide, 2009). A key characteristic of CECs is the presence of multiple fenestrations, allowing for directional flow of oxygen and nutrients and small proteins from the choroid to the RPE (Mrejen and Spaide, 2013), and the removal of waste products from the RPE for systemic recycling. Immunofluorescence and gene expression studies in human AMD donor eyes have demonstrated loss of CECs before RPE degeneration (Mullins et al., 2011, Berenberg et al., 2012, Sohn et al., 2019), resulting in the formation of empty lumens of extracellular matrix named ghost vessels. The outer retina relies heavily on the choriocapillaris for metabolic support; hence, it is thought that the loss of CECs is the trigger for progression to the more advanced stages of the disease.

Several genome-wide association studies have led to the identification of various polymorphisms that affect the risk of developing AMD at all stages. These include single-nucleotide polymorphisms in members and regulators of the complement pathway, including *C3*, *CFI*, *C2* and/or *CFB*, and *CFH* (Fritsche et al., 2014, Tzoumas et al., 2021). The latter is a major negative regulator of the alternative complement system, acting as a cofactor for C3b breakdown and accelerating the decay of C3 convertase (C3bBb), preventing complement hyperactivation (de Córdoba and de Jorge, 2008). One polymorphism in the *CFH* gene, which results in tyrosine (Y) to histidine (H) substitution at amino acid 402 (known as Y402H), increases the risk of AMD by approximately two to sevenfold. Notably, the *CFH* high-risk H allele has been associated with choroid thinning and atrophy. Compared to eyes from donors homozygous for the low-risk Y allele, the choroids of eyes from homozygous donors for the high-risk H allele were 23.6% thinner (Mullins et al., 2014). Furthermore, adults harbouring the *CFH* Y402H polymorphism show increased choroidal C-reactive protein (CRP) (Johnson et al., 2006) and the accumulation of the membrane attack complex (MAC) (Mullins et al., 2011a), which may result in complement-mediated CECs lysis, and may be a primary cause for AMD-associated choriocapillaris degeneration. This upregulation of the complement system was linked to the diminished cell-surface binding activity of the high-risk H402 variant and a corresponding decrease in *in vitro* C3 degradation, as identified through the CFH truncated isoform, FHL-1 (Skerka et al., 2007). Growing evidence suggests a crucial role of AMD-CECs in the development of the pathological environment in the eye, where primary cellular dysfunction is detectable in the choroidal vascular bed.

Most of the work to date characterising the pathology of choriocapillaris in AMD has been performed on post-mortem eyes. While these are very insightful, they have limited availability and represent end-stage disease. We have established a patient-specific induced pluripotent stem cell (iPSC) model of Y402H high-risk AMD RPE (CFH H/H) (Hallam et al., 2017). We have demonstrated that iPSC-derived RPE cells exhibit key cellular features of the disease, including increased inflammation, cell stress, accumulation of lipid droplets, formation of sub-RPE “drusen-like” deposits, complement-mediated reduction in lysosomal function, and impaired waste disposal (Hallam et al., 2017; Cerniauskas et al., 2020). Interestingly, our and other groups’ findings point to the disruption in noncanonical functions of CFH as a crucial factor in AMD pathology, manifesting in the imbalanced mitochondrial homeostasis exhibited in mitochondrial shape shift (Hallam et al., 2017), metabolic impairment (Armento et al., 2020, Ebeling et al., 2021, ten Brink et al., 2025) and diminished capacity to manage oxidative stress (Armento et al., 2020, Armento et al., 2025). Ultimately, this patient-specific platform allows us to interrogate the cellular origins of AMD and explore whether the intrinsic pathologies seen in RPE cells extend to other critical lineages, such as CECs and photoreceptors.

In this study, we utilised a patient-specific iPSC model to generate and characterise CECs, employing high-risk Y402H iPSCs alongside their corresponding isogenic controls, which were derived using CRISPR-Cas9 technology. The generated iPSC-derived CECs express specific markers (CD31, CA4, RGCC, PLVAP) and manifest structural characteristics, including fenestrations, caveolae, and transendothelial cell channels. AMD-iPSC CECs exhibit barrier formation ability and form capillary-like structures similar to the control endothelial cells, with no major differences in glycolysis and mitochondrial function. High-risk CECs exhibit a heightened vulnerability to hydroquinone-induced oxidative damage; this is evidenced by their increased cytotoxicity relative to control CECs and a significant enrichment of C5b-9, the membrane attack complex (MAC) that serves as the terminal product of the complement pathway. Bulk RNA-sequencing data suggest that AMD-CECs undergo significant lipid metabolism remodelling, mesenchymal transition marked by the upregulation of collagen pathways, and increased angiogenesis in response to oxidative stress challenge. These findings highlight a profound disruption in the ability of high-risk CECs to mitigate the oxidative burden imposed by lifestyle-related stressors, such as smoking, poor diet, and physical inactivity, thereby establishing a direct link between environmental triggers and the molecular pathways driving AMD progression.

## Results

### Generation and Characterisation of Y402H and Isogenic iPSC-Derived CECs

To derive endothelial cells manifesting distinct choroid endothelial characteristics, we utilised high-risk (HR) Y402H iPSCs generated in our team from skin biopsies of wet AMD patients (Hallam et al., 2017). To eliminate the potential confounding effect of genetic differences in the control cell lines (Low risk; Y402), we generated isogenic controls (**Figure S1**) using CRISPR/Cas9 gene editing for HR Y402H iPSCs available within our group. A mix of gRNA in complex with Cas9, along with a single-stranded oligonucleotide (ssODN), was delivered to the single-cell suspensions, enabling the homology-directed repair mechanism (**Figure S1**). A PCR screen of isogenic controls for off-target sequences identified for the gRNA (**Table S1**) presented negative results, indicating no relevant off-target effects.

Following a published protocol by Mulfaul et al. (2020), we employed a directed, stepwise differentiation approach to generate CECs. This process began with the formation of mesodermal precursors by treating iPSC-derived embryoid bodies (EBs) with BMP-4, activin A, and FGF-2. Subsequently, these EBs were adhered to Matrigel-coated plates and subjected to VEGF and CTGF to complete their specification into CECs. Following selection on day 14 using anti-CD31 magnetic beads, we expanded HR and corresponding isogenic control CECs (**Figure 1A, B**). To confirm the CEC phenotype of the CD31-positive fractions (Voigt et al., 2019, Mulfaul et al., 2020, Voigt et al., 2021, Li et al., 2024), we performed RT-qPCR analysis for the canonical CEC markers *CD31*, *RGCC*, and *PLVAP* (**Figure 1C**), demonstrating their enriched expression.

**Figure 1:**
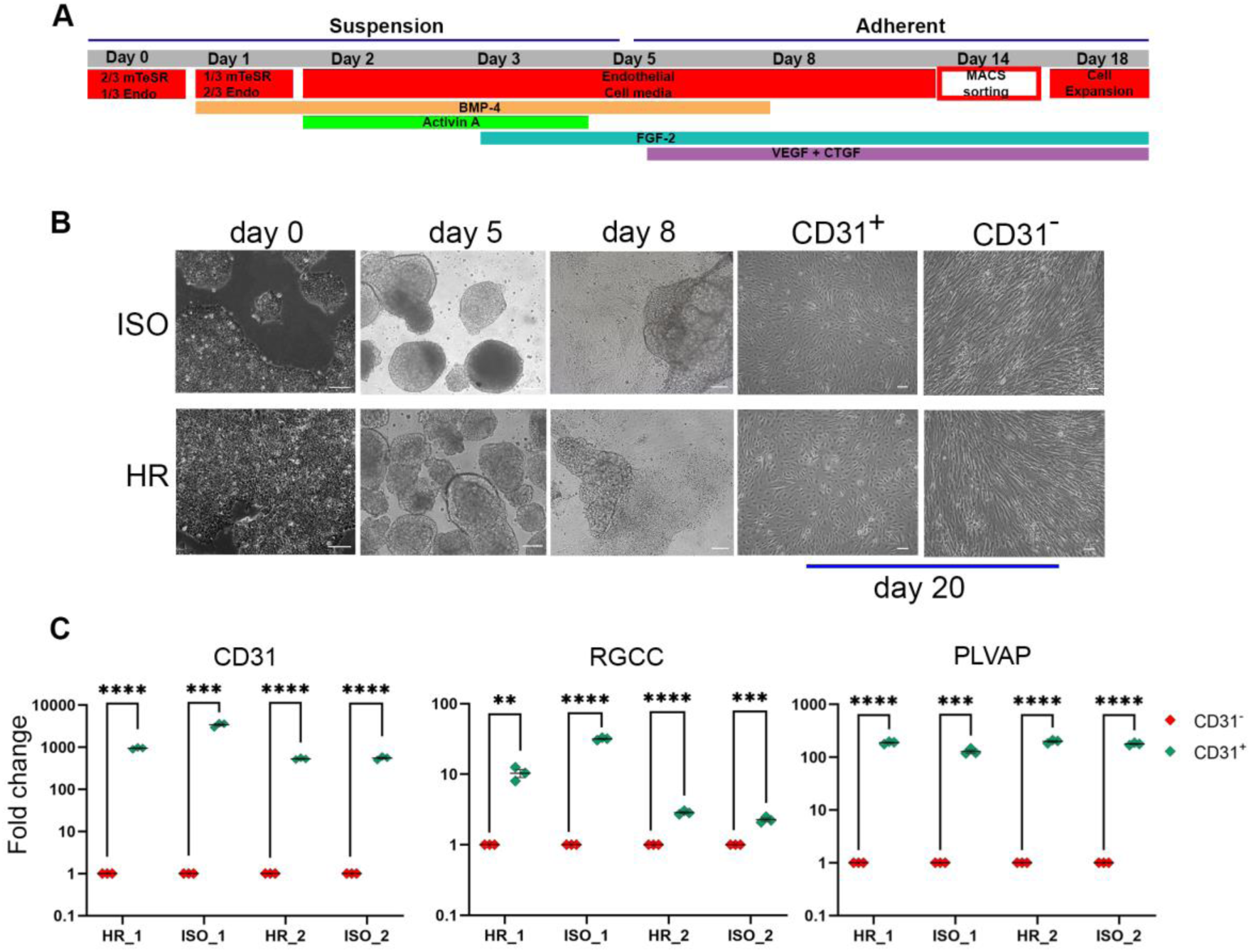
Differentiation of AMD patient-specific iPSCs (HR) and isogenic controls (ISO) to CECs. **A**) Schematic diagram of the stepwise differentiation of iPSCs into CECs. **B**) Bright-field images: day 0; iPSCs, day 5; embryoid bodies, day 8; differentiating cells. The CD31-positive endothelial cells were isolated (day 14) and expanded from the heterogeneous population using the CD31 MicroBead Kit. Representative brightfield images of CD31-negative and CD31-positive cells at day 20. Scale bars = 100 μm. **C**) Quantitative RT-PCR showing enrichment of CEC markers in the CD31^+^ enriched cell populations in both HR and ISO cultures. Data are presented as mean ± SEM, n=3, unpaired t-test. ** p < 0.01, *** p < 0.001, **** p < 0.0001.

To further validate these findings, we performed immunostaining using antibodies previously verified in 12-week post-conception (PCW) human eye sections (**Figure S2**). This analysis confirmed the robust expression of CD31, RGCC, CA4, and PLVAP proteins across both our iPSC-derived HR and ISO CECs (**Figure 2**). The tube formation assay and barrier capacity test were performed to assess the vascular function of iPSC-CECs. The tube formation assay confirmed their ability to form capillary-like structures, visualised by overlaying Calcein-stained images with brightfield images (**Figure S3A**). Their barrier integrity and electrical resistance were comparable to those of control endothelial cell colony-forming cells (ECFCs) (**Figure S3B**). Metabolic profiling of mitochondrial respiration and glycolysis, based on Seahorse Analyser measurements of oxygen consumption rate (OCR) and extracellular acidification rate (ECAR), showed no significant differences in glycolysis or maximal respiration between the iPSC-CECs and ECFCs (**Figure S3C-E**). While ECFCs displayed higher basal OCR values and ATP-linked respiration (**Figure S3D**), it is well-established that endothelial cells derived from distinct vascular beds often exhibit significant metabolic heterogeneity (McAleese et al., 2025).

**Figure 2:**
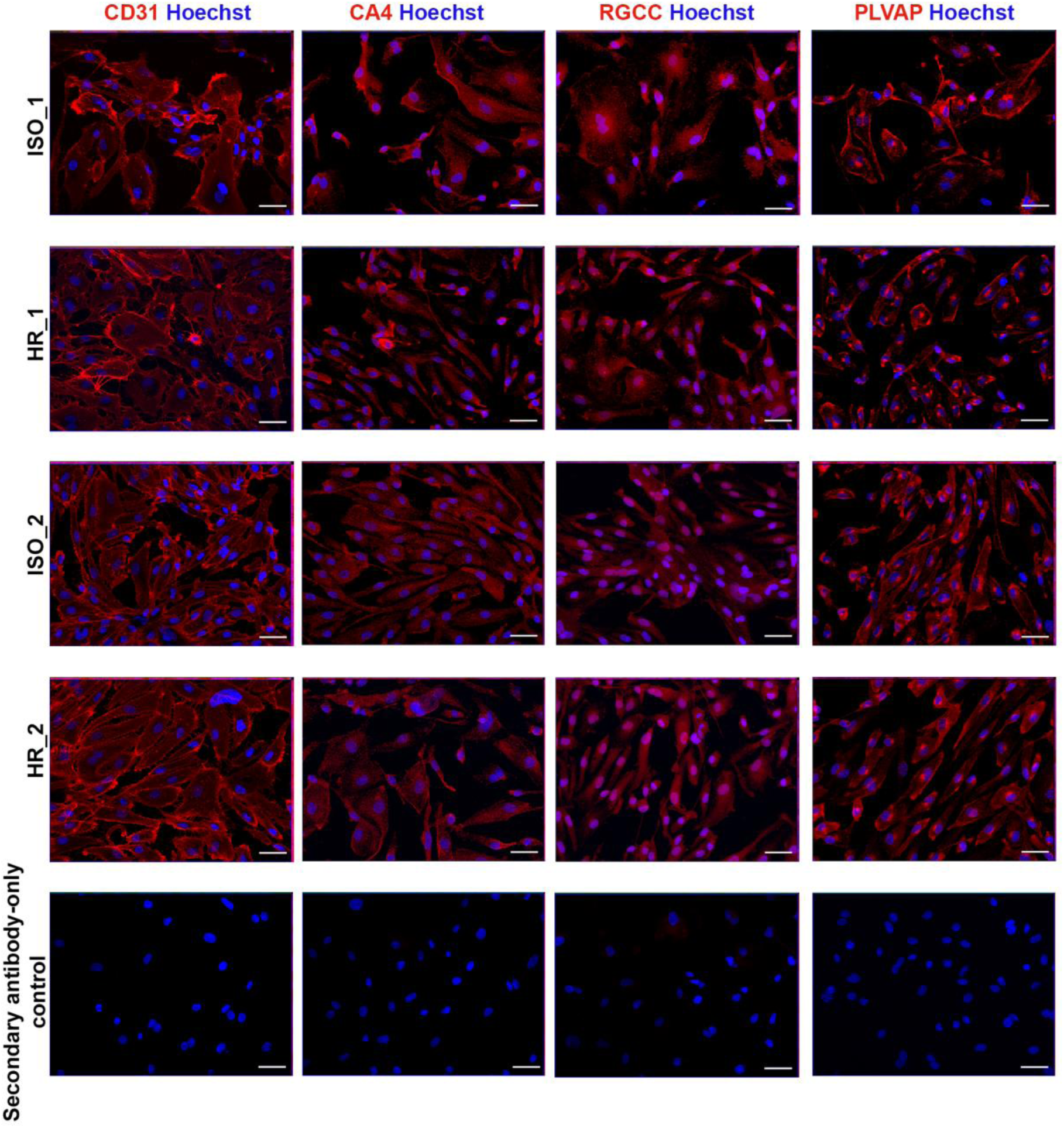
Immunofluorescence analysis of CD31-positive expanded fraction in the high-risk (HR_1 and 2) and iso-control (ISO_1 and 2) derived CECs. CECs were stained for choriocapillaris endothelial cell markers: CD31, RGCC, CA4, PLVAP (red). Nuclei were counterstained with Hoechst (blue). Positive staining was assessed by comparing to a secondary antibody-only control, scale bars = 50 μm.

Further CEC validation encompassed transmission electron microscopy (TEM) assessment of key morphological indicators of maturity. CECs have a unique structure defined by the position and function of the capillary bed in the eye, supplying nutrients and oxygen to the highly metabolically active photoreceptors and removing waste generated by the RPE (Lejoyeux et al., 2022). These features include abundant fenestrations, which facilitate the exchange of water and small molecules like amino acids and sugars, alongside coated pits, caveolae, and vesiculo-transendothelial channels (VECs) that regulate macromolecular transport. Additionally, tight junctions are present to maintain the integrity of the endothelial barrier (Sellheyer and Spitznas, 1988, Sakamoto et al., 1995, Grebe et al., 2019). Both the isogenic and HR CECs successfully recapitulated these structural elements, which are essential for the bidirectional transport of molecules between the systemic circulation and Bruch’s membrane (**Figure 3A-C**).

**Figure 3.**
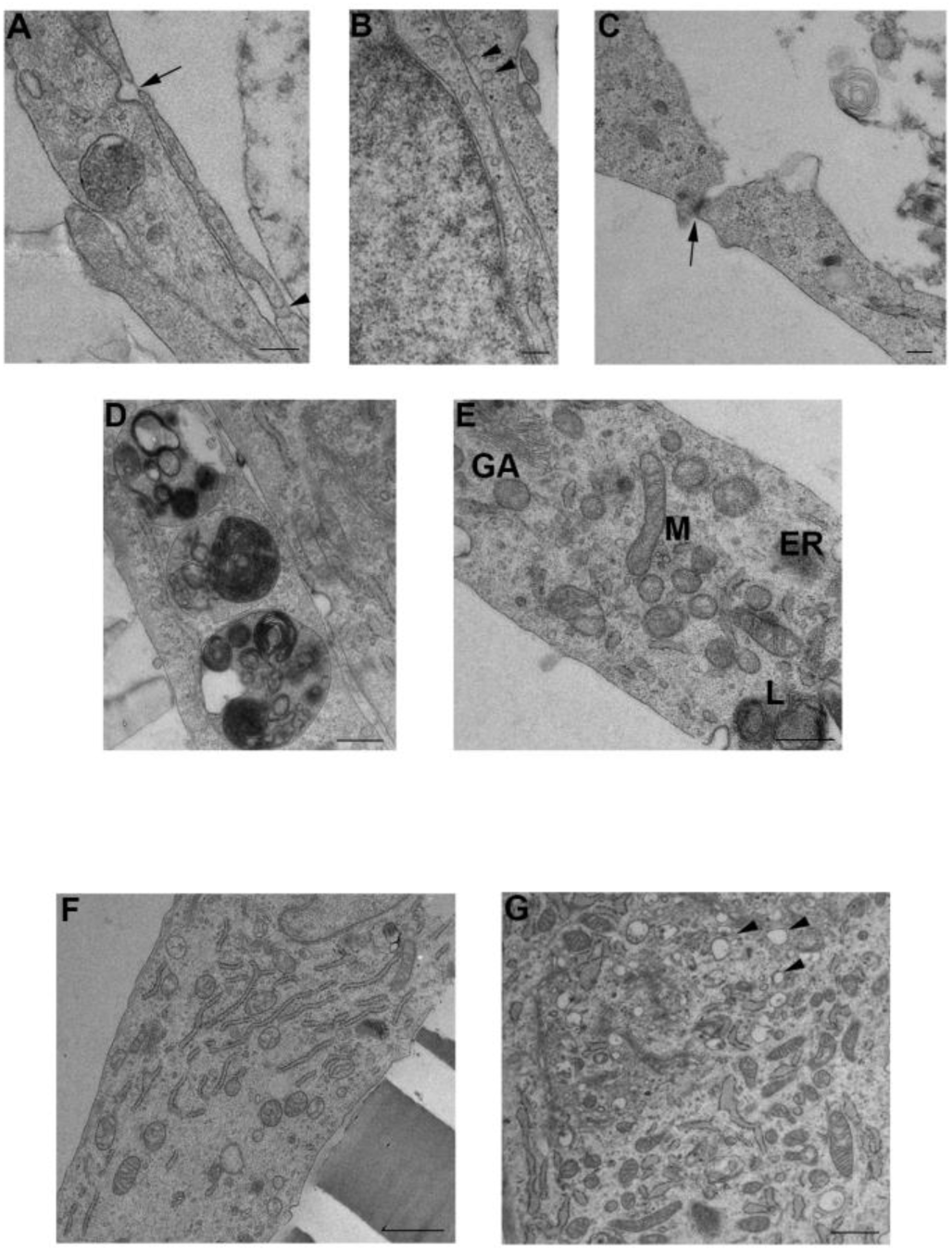
Transmission electron microscopy (TEM) analysis of ISO and HR CECs. **A)** Representative images showing fenestrations (arrow) and transendothelial cell channels (arrowhead). **B)** High-magnification view of caveolae (arrowhead). **C)** Visualization of tight junctions (arrow). Scale bars in A–C: 200 nm. **D)** Representative lysosomes (scale bar: 500 nm). **E)** Subcellular organization in ISO_1 CECs highlighting the Golgi apparatus (GA), mitochondria (M), endoplasmic reticulum (ER), and lysosomes (L) (scale bar: 500 nm). **F, G)** Comparison of ISO_2 and HR_2 CECs; arrowheads in (G) indicate an enrichment of empty vacuoles in HR_2 cells compared to the ISO_2 control (F). Scale bars: 1 μm.

### Complement expression, autophagy, and metabolic flexibility in Y402H and isogenic CECs

The choroid is the major producer of C3 and CFH (Voigt et al., 2021, Zauhar et al., 2022), both of which are critical components at the apex of the alternative complement cascade. In this system, CFH serves as the principal negative regulator of C3 activation. We validated the expression of these regulatory factors, specifically CFH, its truncated isoform FHL1, and CFI, alongside the central components C3 and C5. Bulk RNA sequencing confirmed mRNA expression, though levels varied between the two patient/ ISO-CECs (**Figure S4**). Steady-state intracellular Factor H-like protein 1 (FHL-1) and CFI expression were detected in the HR_1/2 and ISO_1/2 CECs alongside C3, the latter of which is essential for the activation and formation of the MAC (**Figure 4A**). Secreted CFH was confirmed via western blot analysis of CEC supernatants (ISO_1 and HR_1) harvested after a 4-day incubation period (**Figure 4B**). Interestingly, ELISA measurements revealed that only uncleaved C3 was present in the media at very low concentrations; notably, C3 secretion from HR-CECs was significantly lower than that of their ISO counterparts (**Figure 4C**).

**Figure 4.**
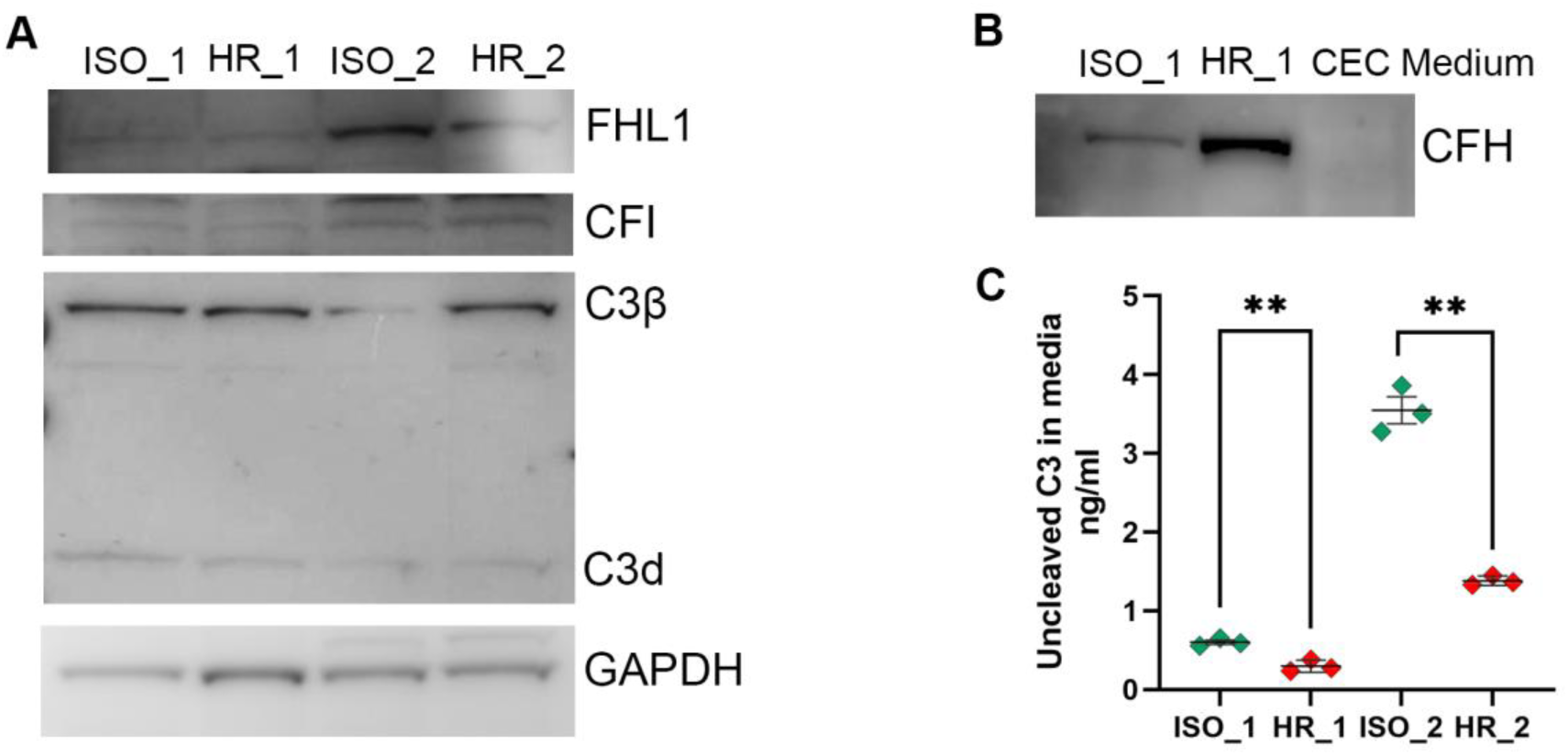
The expression of key complement pathway genes in high-risk (HR) and isogenic controls (ISO) CECs. **A**) Whole cell lysate western blot depicting FHL1, FI, and C3 expression. GAPDH was used as a loading control. **B**) Western blot of secreted CFH by the HR and ISO-CECs. **C**) ELISA analysis of secreted, non-activated C3 in cell culture supernatants of HR and ISO-CECs. Data are presented as mean ± SEM, n=3, unpaired t-test, ** p < 0.01.

A substantial body of research focusing on AMD-RPE cells demonstrates that the high-risk CFH variant fails to perform its regulatory functions. This impairment undermines cellular homeostasis, leading to the dysregulation of phagocytosis, lipid and cholesterol metabolism, and mitochondrial function, while also compromising the cell’s ability to manage oxidative stress (Fisher and Ferrington, 2018, ten Brink et al., 2025, Armento et al., 2025). Building on our previous findings in Y402H iPSC-derived RPE cells, which demonstrated impairments in the autophagy-lysosome pathway (Hallam et al., 2017, Cerniauskas et al., 2020), we evaluated the integrity of the cellular waste disposal and recycling systems within CECs. Western blot analysis revealed no significant differences in the steady-state levels of LAMP1, LAMP2, or p62 across HR-, ISO-, and LR-CECs. A notable exception was the Cathepsin D heavy chain, which showed enrichment specifically in LR-CECs (**Figure S5A**). The autophagic flux (a measure of autophagic degradation activity) in HR-CECs showed no significant difference compared to ISO- or LR-CECs. However, an increase in flux following Bafilomycin A1 treatment was observed in both HR-CEC lines; while this trend did not reach statistical significance, it may serve as an indicator of heightened cellular stress (**Figure S5B, C**). Increased autophagic flux has been documented in the retinas of patients with early-stage AMD, likely as a compensatory response to oxidative stress and intracellular damage (Mitter et al., 2014). Our TEM analysis revealed an enriched presence of empty vacuoles in HR-2 CECs, a hallmark of cellular dysfunction and stress (**Figure 3G**). These membrane-delimited structures have previously been reported in high-risk RPE cells, where they occur alongside mitochondrial remodelling, a process postulated to drive AMD progression (Hallam et al., 2017, Cerniauskas et al., 2020, Kurzawa-Akanbi et al., 2022, Xu et al., 2025, Armento et al., 2025).

To characterise the metabolic signature of AMD-CECs, we performed Seahorse Glycolysis and Mito Stress Tests. These assays were prompted by several studies reporting mitochondrial dysfunction and compromised cellular energy supplies in AMD, characterised by disruptions in both oxidative phosphorylation (OXPHOS) and glycolysis. Maintaining these metabolic pathways is essential for sustaining the balanced ecosystem required for RPE and photoreceptor health (Ferrington et al., 2017, Somasundaran et al., 2020, Ferrington et al., 2020, Ebeling et al., 2021). While CECs are directly exposed to oxygen-rich blood, their primary energy source is glycolysis rather than OXPHOS. This metabolic preference supports their relatively low energy demands in a quiescent state while enabling a rapid surge in energy production during proliferation, migration, and angiogenesis. By relying on glycolysis, CECs preserve available oxygen for diffusion to the underlying retinal layers and ensure vascular function remains uncompromised even in low-oxygen environments (Yang et al., 2021). CECs facilitate the transport of glucose to the mitochondria-rich RPE, which in turn supplies it to the photoreceptors. Notably, photoreceptors utilise glucose via aerobic glycolysis, similar to CECs, and export lactic acid to the RPE. The RPE then utilises this lactate as its primary fuel source through OXPHOS, sparing glucose for the photoreceptors. As some of the most metabolically active cells in the body, photoreceptors rely on a specialised combination of glycolysis and fatty acid oxidation to meet their immense energy demands (Fu et al., 2021). This intricate network of metabolic co-dependencies between the CECs, RPE, and photoreceptors represents a critical frontier in AMD research. Increasingly, metabolic imbalance within the CECs is recognised not merely as a consequence of the disease, but as a primary driver and a vital target for therapeutic intervention (Yang et al., 2021, Chen et al., 2024).

Seahorse metabolic flux analysis showed that both HR and isogenic CECs exhibited low basal ECAR and OCR, followed by a coordinated increase in both glycolytic and respiratory activity after sequential stress challenge. ECAR increased rapidly after the first perturbation and reached a sustained plateau in the stressed phase, indicating enhanced glycolytic engagement. Similarly, OCR rose substantially under stressed conditions as expected, demonstrating preserved mitochondrial respiratory capacity and a strong bioenergetic response to metabolic stress (**Figure S6A, B**). When basal and stressed phenotypes were plotted in ECAR–OCR space, the basal state clustered near a relatively quiescent metabolic zone, whereas stressed conditions shifted the samples toward a more energetic phenotype with higher glycolytic and oxidative activity (**Figure S6C**). Collectively, these data indicate that the iPSC-CECs are metabolically flexible and respond to stress by upregulating both glycolysis and OXPHOS rather than relying on a single pathway. Herein, ISO_1 CECs shifted further upward and rightward compared to HR_1 CECs, implying stronger activation of both respiration and glycolysis. Biologically, that pattern is consistent with differences in metabolic resilience or stress adaptability, rather than major baseline metabolic differences (Epstein et al., 2014).

Together, these data show that iPSC-derived CECs express key alternative complement components and regulatory factors, supporting the concept of local complement production and control within the choroidal endothelium. Importantly, although CFH and related transcripts and proteins were detectable in CECs, C3 abundance remained low, consistent with the idea that CEC-derived complement output is modest compared with other ocular cell types such as RPE cells. This low-level C3 secretion, together with intact autophagic flux and a preserved ability to increase both glycolytic and respiratory metabolism under stress, suggests that iPSC-CECs maintain homeostatic resilience under baseline conditions. However, the increased stress-responsive metabolic shift observed in ISO_1 CECs indicates that AMD-associated cells may exhibit altered bioenergetic adaptability and stress handling, which could contribute to endothelial dysfunction in disease.

### High-risk CECs manifest higher sensitivity to oxidative stress

To assess the role of complement activation in CEC loss in early AMD, we treated high-risk CECs and their isogenic controls with human serum and its inactivated form, as well as RPE conditioned medium. Both human serum and high-risk RPE conditioned medium were shown to be potent inducers of the AMD phenotype in RPE cells (Sharma et al., 2021, Kurzawa-Akanbi et al., 2022). The IncuCyte cytotoxicity assay was utilised to assess the viability of high-risk and isogenic CECs exposed to 5% human serum and its inactivated form (**Figure 5A**) or conditioned medium harvested from HR- and LR-RPEs (**Figure 5B**). Cell death was assessed by measuring the uptake of fluorescent dyes into cells with compromised membrane integrity. This allowed for the quantification of dying cells and provided a robust measure of cytotoxicity, which was further validated by confluency analyses. The addition of either human serum (HS) or heat inactivated human serum (IHS) increased cytotoxicity in both HR and isogenic CECs for pair 1. However, across both pairs, no significant differences in confluency (data not shown) or cytotoxicity were observed between the HR and isogenic-derived cells (**Figure 5A**). Similar results were obtained for exposure to 50% HR and LR RPE conditioned medium (**Figure 5B**). These results suggest no direct effect of extracellular complement system activation on high-risk CECs.

**Figure 5.**
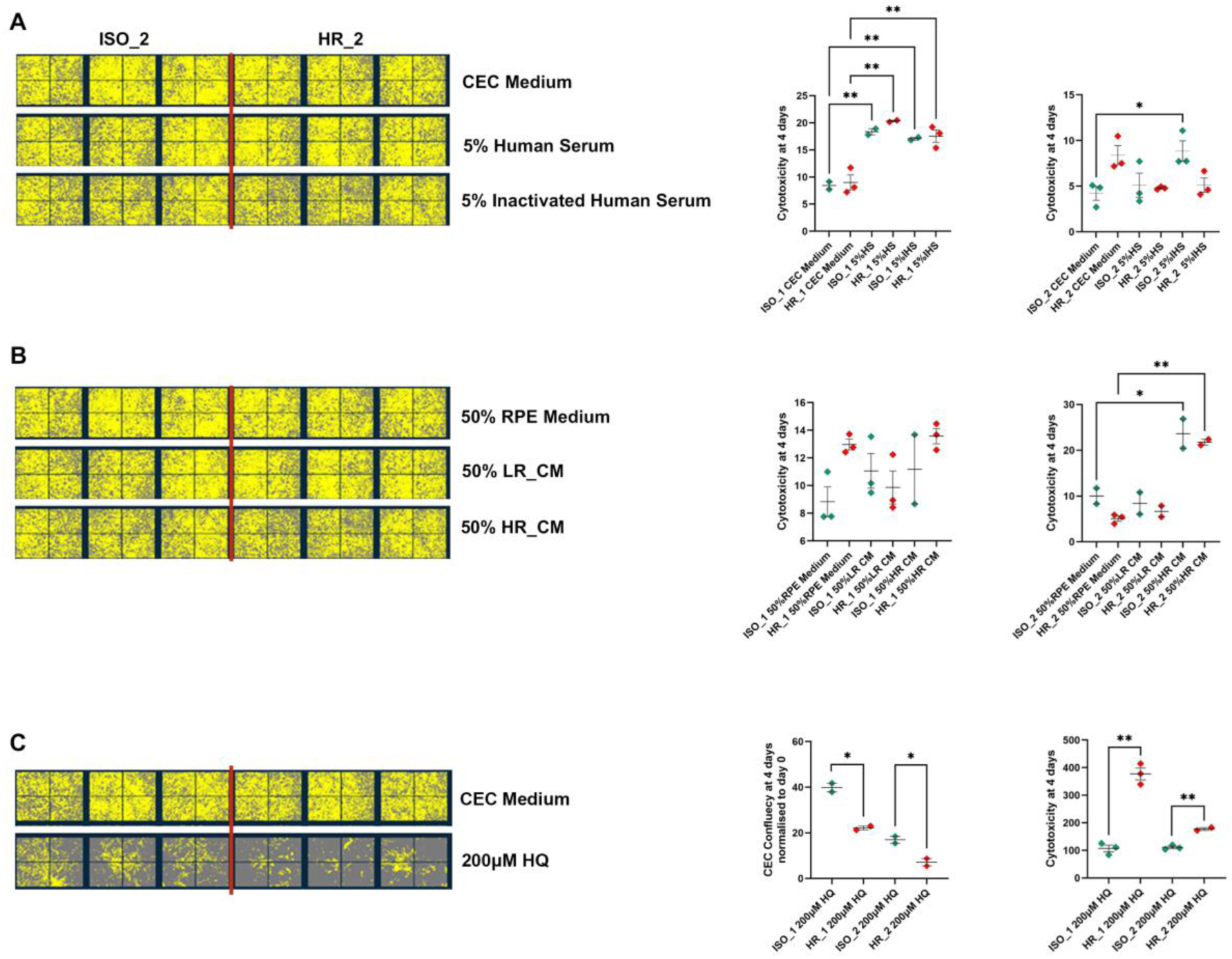
Hydroquinone (HQ) induces higher cytotoxicity in high-risk (HR) CECs in contrast to human serum and RPE high-risk conditioned medium after 96-hour exposure. **A**) Representative images of CEC high-risk (HR_2) confluency in the presence of human serum (HS) and inactivated human serum (IHS) versus isogenic control (ISO_2). **B**) The growth conditions were changed to 50% RPE medium (control condition), low-risk conditioned medium (LR_CM), and high-risk conditioned medium (HR_CM). **C**) Images present the HR_2 CECs confluency in the presence of Hydroquinone versus the isogenic control. **A-C**: The graphs on the right show quantified cell cytotoxicity and confluency (C). Data are presented as mean ± SEM (n =3), two-way ANOVA, * p < 0.05, ** p < 0.01.

Smoking is a major modifier of AMD, increasing the risk of both developing and worsening the existing condition (Velilla et al., 2013). Human RPE cell line treated with cigarette smoke extract manifested increased mitochondrial reactive oxygen species level, altered mitochondrial morphology and downregulation of glycolytic and mitochondrial ATP production (Henning et al., 2023). Recent studies demonstrate that high-risk RPE cells exposed to hydroquinone (HQ), a pro-oxidant component of cigarette smoke, exhibit diminished resilience to reactive oxygen species (ROS) following a 48-hour exposure (Armento et al., 2025). To assess the role of oxidative stress on the CECs, we exposed HR and ISO CECs to 200 μM Hydroquinone (HQ) for 96 hours after confirming heightened levels of hydrogen peroxide (H_2_O_2_) in its presence (**Figure S7**). HR-CECs showed higher sensitivity to HQ, demonstrated by significantly lower confluency and increased cytotoxicity in comparison to ISO-CECs (**Figure 5C**). Immunocytochemical staining of CECs after 24 hours of exposure to 200 µM HQ presented a striking difference in expression of membrane attack complex (C5b-9) between HR-CEC_2 and ISO_2, with highly enriched detection of C5b-9 in AMD-CECs and augmented expression of CASP3 (**Figure 6A** and **B**). Functional test assessing the tube formation ability of HR_2 in comparison to ISO_2 CECs showed that both groups formed a significant network with thick endothelial cell wall forming vessel-like lumen structures. The presence of HQ lowered vascular density, vessel length density, and vessel thickness of HR-CECs (**Figure S8**).

**Figure 6.**
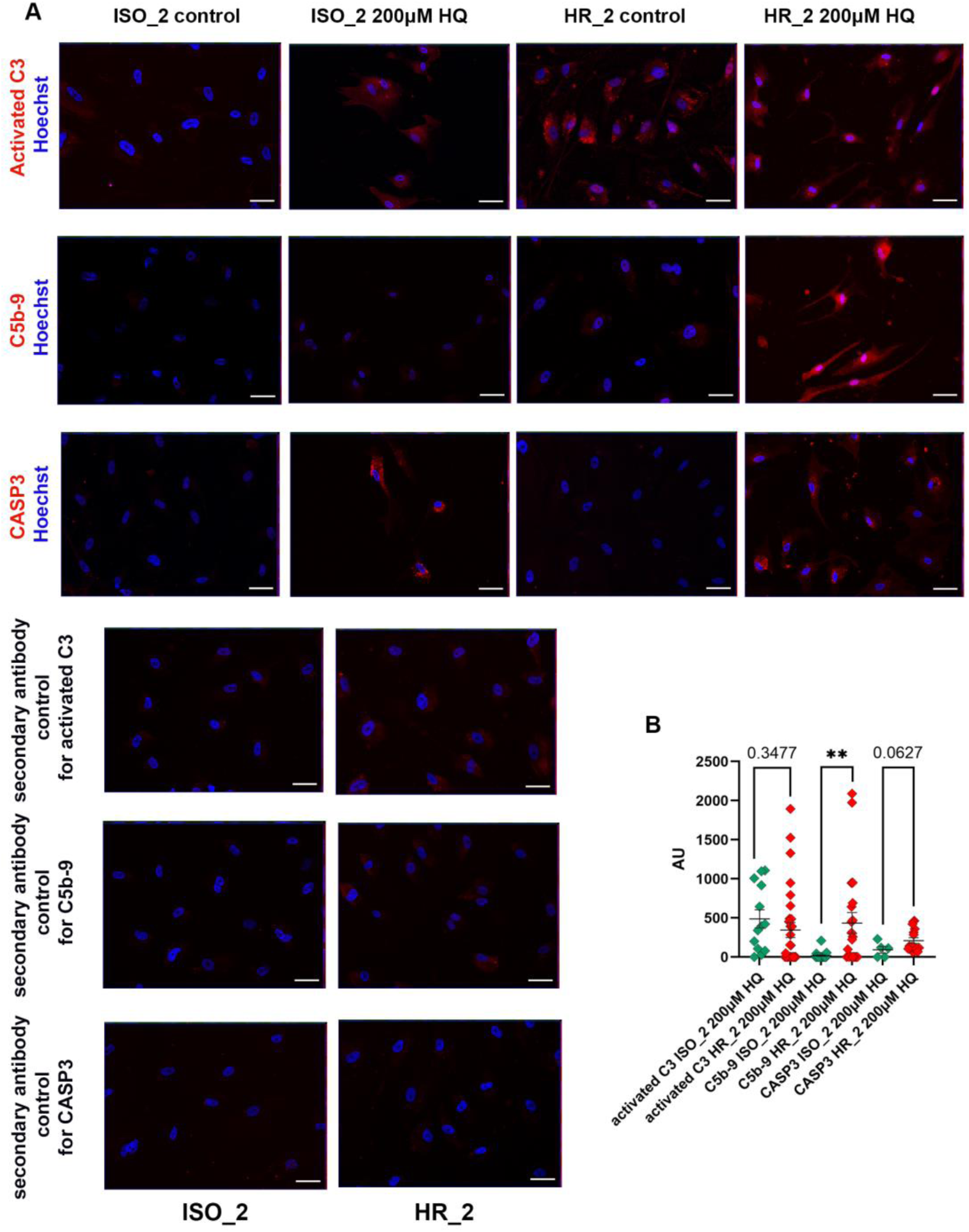
High-risk CECs display an enriched level of C5b-9 and activated Caspase 3 in the presence of 200 μM HQ. **A**) Representative immunofluorescent staining of activated C3, C5b-9, and CASP3 counterstained with Hoechst. Scale bars: 50 µm. **B**) Graphs showing intensity of fluorescence from activated C3, C5b-9, and CASP3 in the presence of 200 μM HQ, acquired with Image J analysis. Data are presented as mean ± SEM (n = 3), unpaired t-test, ** p < 0.01.

### Oxidative Stress–Induced Remodelling of Lipid Metabolism, Fibrosis, and Angiogenesis in CECs

To further investigate the response of high-risk CECs to the presence of hydroquinone relative to their ISOs, we next performed bulk RNA-Seq on both pairs subjected to 200 μM HQ for 6 hours (**Figure 7A**). The PCA reveals a clear separation between HR and ISO CECs along PC1 for patient 1 with a less clear separation for patient 2, indicating distinct global transcriptional profiles between the two groups (**Figure 7B**). Within each pair, HQ treatment causes additional separation from untreated controls along PC2 (24% variance), suggesting that HQ induces significant gene expression changes in both HR and ISO CECs. A substantially higher number of differentially expressed genes (DEGs) was identified for HR_1 versus ISO_1 CEC comparison than for HR_2 and ISO_2 (**Table S4**) under baseline (1408/238) and HQ treated (2446/263) conditions, respectively.

**Figure 7.**
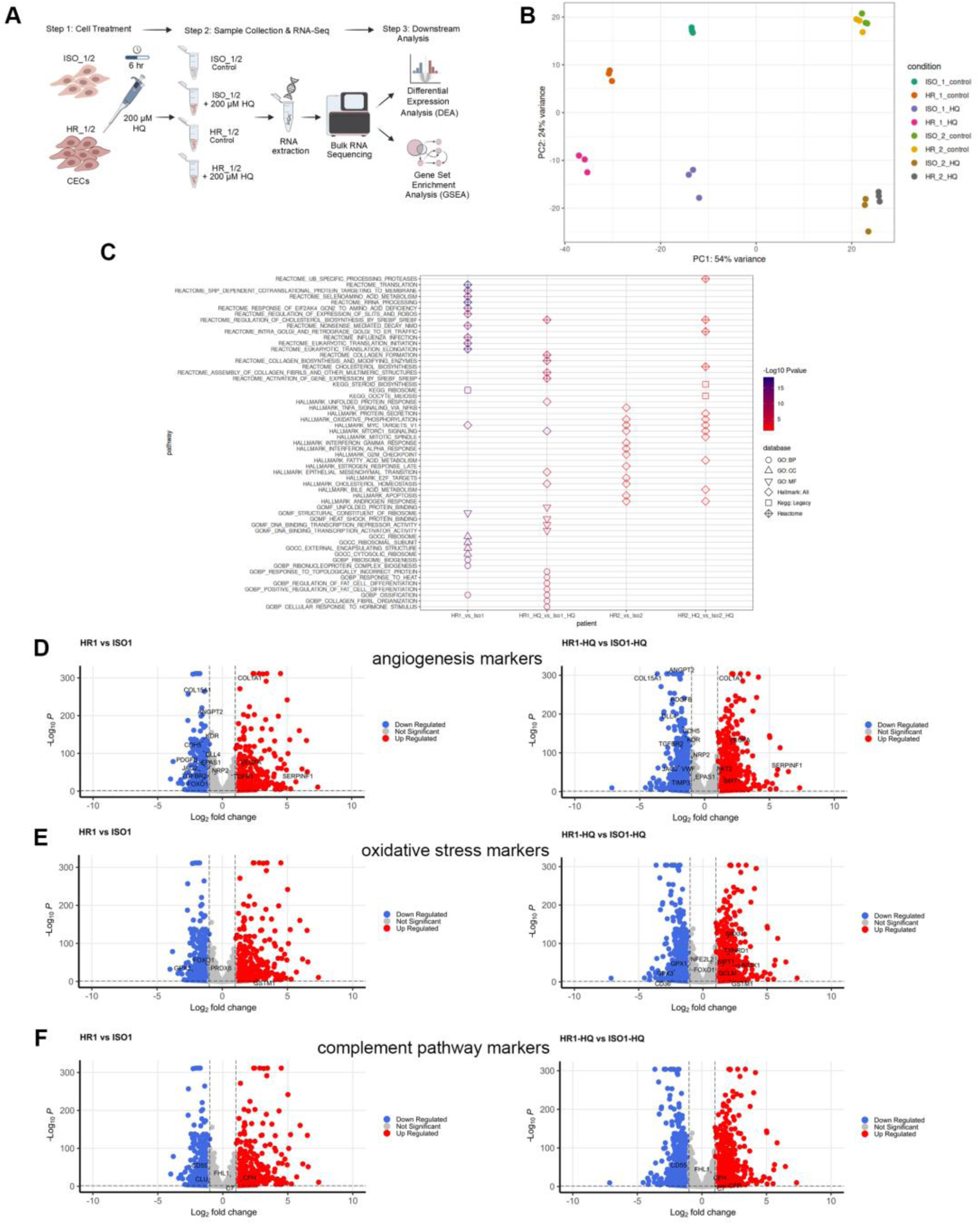
Bulk RNA sequencing analyses depict augmented lipid metabolism, endothelial to mesenchymal transition and angiogenesis remodelling in HR-CECs under oxidative stress. **A**) Schematic representation of the experimental approach for bulk RNA sequencing analyses. **B**) Principal component analysis depicting transcriptional variations between samples. **C**) GO, Hallmark, KEGG and Reactome analyses of upregulated DEGs in HR_1/2 CECs in comparison to ISO_1/2 in the presence of HQ/control conditions. The top twenty pathways for each contrast were selected for visualisation based on p-value. Volcano plots depicting angiogenesis (**D**), oxidative stress (**E**) and complement pathway (**F**) marker gene expression in HR_1 compared to ISO_1 cells under HQ and control conditions. DEG: differentially expressed gene. DEGs were identified from relevant contrasts using DESeq2. Significance thresholds shown, absolute Log_2_ fold change of 1 and adjusted p-value < 0.05.

Pathway enrichment analysis for the top 20 HR_1/2 versus ISO_1/2 listed pathways in control and HQ conditions identified 19 pathways with upregulated gene expression uniquely associated with HQ exposure in HR_1 and 10 in HR_2 CECs compared to their isogenic controls (**Figure 7C, Table S5**). In addition, 1 pathway was shared between HQ and control conditions in HR_1 and 4 in HR_2, while 19 and 8 pathways were specific to control conditions in HR_1 and HR_2 versus isogenic CECs, respectively. In contrast, 18 pathways were identified with significantly downregulated genes in HQ-treated HR_1 versus isogenic CECs, while 2 pathways were shared between HQ and control conditions, and 4 were specific to the control condition (**Figure S9A** and **Table S5**). No enriched pathways were identified for downregulated DEG genes in HR_2 versus isogenic CEC control under HQ treatment and baseline conditions. Together, these data suggest that HQ treatment reshapes pathway activity in HR CECs, with distinct and patient -specific enrichment patterns rather than a single uniform response.

The response of both high-risk CEC lines in the presence of ROS included cholesterol biosynthesis by SREBP upregulation and mTORC1 signalling (**Figure 7C**). Upregulation of steroid biosynthesis and fatty acid metabolism pathways was also noted in HR_2 CECs when compared to their isogenic controls. ROS can promote *de novo* cholesterol synthesis by increasing glucose consumption and activating SREBP2-dependent transcription, including LDLR and caveolin-1, which enhances exogenous cholesterol uptake and may contribute to pathological lipid accumulation (Seo et al., 2019). Upregulation of steroid biosynthesis and fatty acid metabolism genes may initially serve as an adaptive response to oxidative stress and mitochondrial maintenance, but sustained activation can drive lipid accumulation and chronic pathology (Li et al., 2025, Yoon et al., 2021). Accumulation of esterified cholesterol and apolipoprotein-containing lipid particles within Bruch’s membrane and drusen is a recognised feature of AMD pathology and may reflect dysregulated lipid synthesis, export, and retention in the ageing outer retina (Wang et al., 2010, Curcio et al., 2011). Saturated fatty acids are considered lipotoxic and can trigger apoptosis, inflammation, ceramide production, ROS generation, nucleolar stress, and ER stress (Yoon et al., 2021). Growing evidence suggests that aberrant lipid metabolism, extracellular lipid deposition and complement dysregulation are linked in AMD. Lipid abnormalities, including disturbed cholesterol homeostasis, oxidised lipid accumulation, and lipofuscin deposition, have been implicated in complement activation (Tang et al., 2024). Consistent with this, a Chinese case-control study of patients with neovascular AMD and polypoidal choroidal vasculopathy reported a significant interaction between *rs3764261* in *CETP* and *rs800292* in *CFH*, for both disease subtypes (Liu et al., 2014).

Across the HQ conditions, we also observed shared mTORC1 signalling, suggesting a common stress-adaptive response in both models. In the retina, mTOR pathways are closely tied to nutrient sensing, survival signalling, and metabolic reprogramming, all of which are relevant to AMD pathobiology. This is particularly important in the context of oxidative stress, which is a central driver of RPE dysfunction in AMD. mTOR signalling can influence how cells respond to oxidative damage by regulating protein synthesis, autophagy, and energy balance, and prolonged dysregulation may shift cells from adaptation toward degeneration. In parallel, mTOR activity is also linked to cholesterol and lipid biosynthesis, so its shared activation under HQ conditions may reflect an attempt to restore membrane homeostasis and metabolic stability, but one that could also contribute to lipid accumulation when sustained (Huang et al., 2019, Go et al., 2020, Wang et al., 2022).

HQ exposure in HR_1 CECs upregulated epithelial–mesenchymal transition–associated genes relative to isogenic CECs (**Figure 7C**). This is consistent with reports that oxidative stress can drive mesenchymal and profibrotic programs in AMD models, including increased extracellular matrix deposition and COL1A1 expression (Wang et al., 2026). In parallel, collagen formation and fibril organisation pathways were enriched, while endothelium development pathways were downregulated, supporting a shift toward a more fibrotic, less endothelial phenotype (**Figure 7C** and **Figure S9A**). In early AMD, CEC apoptosis is a prominent feature, whereas in wet AMD, surviving CECs can contribute to angiogenesis through VEGFA upregulation and dysregulated DLL4–Notch signalling, promoting excessive sprouting, vascular disorganisation, and ultimately choroidal neovascularisation (CNV) (Jiang et al., 2025; Suchting et al., 2007). Consistent with this, analysis of angiogenesis-related DEGs in HR_1 showed downregulation of *COL15A1, ANGPT2, KDR, CDH5, PDGFB, EPAS1, JAG2, TGFBR2, FOXO*, and *DLL4*, alongside upregulation of *COL1A1*, *VEGFA*, and *SERPINF1* in untreated cells, with HQ further modifying this profile (**Figure 7D**). Notably, HQ further downregulated *ANGPT2 (*2.5 log fold change relative to untreated baseline comparison), *PDGFB* (2.4), *COL15A1*(2), *TGFBR2* (1.6), and *DLL4* (1.4) expression, while *COL1A1* remained elevated, *VEGFA* was further increased (1.4), and *SIRT1* was specifically upregulated in the HQ condition. These changes point to impaired vascular stabilisation and a shift toward a pro-angiogenic, unstable endothelial phenotype. Reduced*, PDGFB, COL15A1, TGFBR2,* and *DLL4* may weaken pericyte support, barrier integrity, and TGF-β/Notch-dependent vessel maintenance, while *SIRT1* upregulation may reflect a stress-adaptive but potentially pro-angiogenic response in the HQ condition (Lin et al., 2024, Schlecht et al., 2017, Eklund et al., 2001, Maloney et al., 2013, Balaiya et al., 2012). Interestingly, HR_1 CECs also upregulated *SERPINF1*, which encodes pigment epithelium-derived factor (PEDF), an anti-angiogenic protein that can suppress VEGF signalling and has been explored as a therapeutic target in choroidal neovascularisation (Jakobsen et al., 2024, Cabral et al., 2017). However, *SERPINF1*, while still overexpressed in HR_1 compared to isogenic CECs, decreased 1.9-fold after 6 hours of HQ treatment compared to baseline treatment, suggesting that oxidative stress may weaken this anti-angiogenic counterbalance. This is consistent with reports of reduced PEDF expression in AMD RPE and after repetitive HQ-induced oxidative injury in ARPE-19 cells (Pons and Marin-Castaño, 2011).

HQ exposure was associated with a coordinated oxidative-stress response in HR_1 CECs, marked by reduced *CD36* and *GPX1* and increased *GCLM, HMOX1, SIRT1*, *TXNRD1, SRXN1,* and *TXNRD2* expression in HR_2 CECs (**Figure 7E and Figure S9C**). This pattern is consistent with a shift away from CD36-mediated anti-angiogenic and lipid-clearance functions and toward activation of antioxidant defence pathways, which may reflect an adaptive response to HQ-induced redox stress (Silverstein and Febbraio, 2009, Febbraio et al., 2001, Yang et al., 2002, Castilho et al., 2012, Lee et al., 2019, He et al., 2021, Wang et al., 2025, Go and Jones, 2008, Cáceres-Vélez et al., 2022).

HQ exposure shifted the complement profile toward dysregulation, with *CFP* (a positive regulator of the complement alternative pathway) upregulated in HR_1 and *CFH* upregulated in both control and HQ conditions, while *CD55* remained downregulated in HR_1 (**Figure 7F**). *FHL1* upregulation was characteristic of HR_2 only in control samples (**Figure S9D**). This suggests increased alternative pathway amplification alongside reduced membrane protection, a combination that would favour complement activation and is consistent with AMD-associated inflammatory stress (Kouser et al., 2013, Blatt et al., 2016, Sweigard et al., 2014, Yue et al., 2025).

HQ exposure also altered programmed cell death genes, with *MCL1* (a critical anti-apoptotic member of the BCL-2 family that promotes cell survival) upregulated in HR_1, while *CASP10* and *TNFRSF10A* were downregulated in control conditions (**Figure S9F)**; no significant changes were detected in HR_2 (Sancho et al., 2022, Kumari et al., 2019, Mori et al., 2022). Together, these changes suggest a shift toward enhanced cell survival and reduced apoptotic signalling under HQ stress.

Six hours of HQ treatment did not produce a consistent inflammatory response in high-risk CECs **(Figure S4**). *TNFα* was strongly upregulated in HR_1, whereas in HR_2 it increased only modestly and remained close to ISO levels. *FHL1* was elevated in both HR lines, indicating a partial but heterogeneous stress response, similar to the cell line-dependent inflammatory effects reported in AMD-RPE models carrying rare CFH variants (ten Brink et al., 2025).

## Discussion

Histopathological analyses reveal that choriocapillaris loss is an early event in AMD, occurring before RPE and photoreceptor degeneration (Mullins et al., 2011, Sohn et al., 2019, Mulfaul et al., 2022). An intertwined symbiotic relationship between photoreceptors, RPE, Bruch’s membrane and choriocapillaris network, where the latter is the sole provider of oxygen and nutrients and waste removal route in the macula, strongly implicates vascular insufficiency in AMD pathophysiology. Given the vital role of choriocapillaris in maintaining the homeostasis of the outer retina and its early diminished presence in AMD eyes, we focused our research on choriocapillaris endothelial cells. We successfully generated and characterised CECs from two iPSC lines carrying the *CFH* Y402H high-risk polymorphism, alongside CRISPR-Cas9–engineered isogenic controls; this robust framework ensures that any observed phenotypes can be directly attributed to the high-risk variant rather than inter-individual genetic variability. The differentiated cells expressed canonical CEC markers (CD31, RGCC, CA4 and PLVAP: Voigt et al., 2021), formed capillary-like networks and exhibited barrier formation capability similar to the unaffected control CECs. Moreso, iPSC-derived CECs displayed ultrastructural hallmarks of mature choriocapillaris, including fenestrations, caveolae, transendothelial cell channels and tight junctions. These characteristics define the critical role of the choriocapillaris endothelial cells in the specialised transport requirements at the blood retinal barrier, where the precise balance between high metabolic demand and the immune-privileged environment of the retina must be upheld for its integrity and healthy function.

High-risk CECs did not manifest inborn defects in glycolysis or mitochondrial respiration compared with isogenic controls. AMD is a multifactorial disease, developing over a prolonged period of time, hence the metabolic fitness of CECs may not be defined by genetic risk alone. Even though mitochondria are not the main source of ATP for endothelial cells, they play crucial regulatory roles in redox balance, calcium handling, and apoptosis. Stressors like smoking, heightened alcohol intake, and the metabolically demanding environment of the macula, combined with genetic factors, could therefore lower the threshold for dysfunction. Local *CFH, FHL1*, and *CFI* synthesis at the mRNA level was confirmed in high-risk and control CECs, although *CFH* and *CFI* transcript expression was diminished in HR_2 and ISO_2. At the protein level, CFH short isoform, FHL-1, was detected at low expression levels in both CEC pairs alongside CFI and C3, secreted CFH was also confirmed in CEC supernatants; these findings support the presence of localised, albeit modest, complement production. Interestingly, high-risk CECs secreted lower levels of C3 compared with isogenic controls, upholding the same pattern as observed in our AMD-RPE analyses. However, the overall level of detected uncleaved C3 was far higher in RPE medium (Cerniauskas et al., 2020). The largely lower level of C3 secretion by CECs is probably the reason for the failure to detect C3 fragments in CEC medium (for ISO_1 and ISO 2: ∼500 and 90 times respectively, HR_1 and HR_2: 160 and 40 compared to RPE counterparts).

Endothelial autophagy is inherently cytoprotective, regulating the function of endothelial cells in response to blood flow and stress. Moreover, it affects several aspects of endothelial cell characteristics, including sprouting, permeability and secretion (Mameli et al., 2022, Hu et al., 2024). In CECs, we observed preserved expression of autophagy markers; LAMP1/2, Cathepsin D and p62 across high-risk and isogenic controls. Autophagic flux did not present significant changes in LC3_II expression, although a pattern of increased bafilomycin A1–induced flux, especially for HR_2 CECs, was noticed. The latter, along with the presence of empty vacuoles on TEM, may suggest heightened cellular stress in high-risk cells and possibly an early adaptive response.

A key finding of this study is that high-risk Y402H CECs are more vulnerable to oxidative stress induced by hydroquinone, a major component of cigarette smoke and a well-established AMD risk modifier (Sharma et al., 2012, Armento et al., 2025). Exposure to hydroquinone caused a greater reduction in confluency and higher cytotoxicity in high-risk CECs than in isogenic controls. AMD CECs in the presence of HQ showed significant enrichment of C5b-9 deposition, alongside increased CASP3 expression. Notably, this vulnerability was not reproduced by exposure to 5% human serum or 50% high-risk RPE-conditioned medium, suggesting that extracellular complement activation alone is insufficient to drive AMD CEC death. Instead, these data support a model in which oxidative stress triggers a defective complement response in Y402H CECs, leading to significant MAC formation not detected in isogenic control CECs. This dependence between oxidative stress and the high-risk Y402H polymorphism, manifesting as MAC-mediated injury in AMD CECs, aligns well with the multifactorial nature of AMD (Armento et al., 2021, Saigal et al., 2025). The absence of a strong phenotype under basal conditions supports a multifactorial trigger that favours environmental insult in combination with genetic predisposition.

Bulk RNA-Seq analysis revealed a pronounced differential gene expression response in HR_1 and HR_2 CEC lines exposed to HQ, with HR_1 manifesting a far broader and more complex transcriptional landscape. Interestingly, both high-risk lines converge on lipid metabolic reprogramming in response to HQ-induced ROS. In particular, upregulation of cholesterol biosynthesis pathways in both high-risk cell lines suggests a shift toward enhanced de novo lipid synthesis and uptake. While such adaptations may initially support membrane repair and cellular survival under oxidative stress, sustained activation comes at the cost of long-term cellular health, promoting lipid accumulation, mitochondrial dysfunction, and chronic cellular stress.

The observed upregulation of steroid biosynthesis and fatty acid metabolism pathways in HR_2 further supports a model in which metabolic remodelling serves as a compensatory mechanism to preserve mitochondrial integrity. However, accumulation of saturated fatty acids and lipid intermediates can exacerbate oxidative damage, induce endoplasmic reticulum stress, and activate inflammatory signalling cascades. These findings align with growing evidence that lipid metabolism dysregulation is an active driver of AMD progression, involving its crosstalk with the complement system (Chen et al., 2024, Tang et al., 2024). The specific complement profile obtained in our study, characterised by *CFP* and *TNFα* upregulation alongside the downregulation of the regulator *CD55*, indicates that metabolic remodelling in high-risk CECs likely feeds into a pro-inflammatory feedback loop with the complement system, mirroring the signature seen in clinical AMD (Tang et al., 2024).

A striking feature of the HQ response in HR_1 cells is the activation of mesenchymal and collagen-associated pathways, indicative of endothelial-to-mesenchymal transition (EndMT). Concurrent downregulation of the endothelial development pathway and endothelial marker *VWF* further supports this transition, alongside persistently upregulated *COL1A1* in both HQ and control conditions. EndMT has been increasingly implicated in vascular dysfunction and fibrosis in neovascular AMD, where senescent, ROS-exposed endothelial cells contribute to excessive ECM deposition and loss of vascular integrity (Chirco et al., 2017, Wang et al., 2026). HQ also induces complex modulation of angiogenic signalling in HR_1, revealing simultaneous upregulation of pro-angiogenic factors such as *VEGFA* and *SIRT1*, alongside downregulation of key regulators including *ANGPT2, PDGFB, TGFBR2*, and *DLL4*. This pattern suggests a state of angiogenic dysregulation favouring neovascularisation. Increased *VEGFA* expression is consistent with ROS-driven angiogenesis observed in wet AMD (Zuo et al., 2025), while SIRT1 upregulation may reflect an adaptive attempt to counteract oxidative stress through mitochondrial biogenesis and antioxidant activation (Campagna et al., 2024). However, given evidence linking SIRT1 to enhanced VEGF signalling and choroidal neovascularisation, its role may be context-dependent and potentially pathogenic under sustained stress (Maloney et al., 2013). Conversely, the downregulation of *PDGFB* and *TGFBR2* is likely to compromise vascular stability by impairing pericyte recruitment and TGF-β-mediated homeostatic signalling, respectively (Lin et al., 2024, Schlecht et al., 2017). Similarly, reduced expression of DLL4 and other Notch pathway components may contribute to excessive, disorganised vascular sprouting (Suchting et al., 2007, Jiang et al., 2025).

The oxidative stress response in HR_1 is characterised by induction of antioxidant defence systems, including upregulation of *GCLM, HMOX1, TXNRD1, SRXN1*, and in HR_2 mitochondrial *TXNRD2*. These changes reflect activation of cellular redox buffering mechanisms aimed at restoring homeostasis. However, the simultaneous downregulation of *GPX1, GPX3* and *CD36* in HQ-treated HR_1 suggests that these compensatory responses are incomplete or imbalanced, potentially leading to choriocapillaris endothelial bed damage and development of neovascularisation (Silverstein and Febbraio, 2009, Sharma et al., 2016, Chen et al., 2024). Importantly, prolonged or excessive activation of stress-response pathways such as *HMOX1* may become deleterious, contributing to endoplasmic reticulum stress and apoptosis (Li et al., 2021).

Alterations in programmed cell death pathways support a survival preservation yet dysfunctional phenotype in HR_1 cells. Upregulation of the anti-apoptotic factor *MCL1* in the presence of HQ, and downregulation of pro-apoptotic mediator *CASP10* in control conditions indicate a shift toward apoptosis resistance. While this may enable short-term survival under oxidative stress, it may also permit the persistence of damaged, malfunctioning endothelial cells that contribute to pathological remodelling and angiogenesis.

Some limitations should be acknowledged in our study. First, while this model successfully isolates CEC-intrinsic defects, it inherently lacks the multicellular complexity of the native outer retina, omitting the dynamic reciprocal signalling between CECs and their neighbouring pericytes, fibroblasts, immune cells, and the RPE. It would be beneficial for the CECs to be incorporated into the multilayer model spanning the mentioned above cell types with an active perfusion through the choriocapillaris, a model similar to Song et al., 2023, although the latter lacked the immune cells and high-volume flow. The multicellular environment burdened with high-risk polymorphism would allow us to conduct comprehensive analyses of the codependent development of AMD phenotype and test possibly multifactorial treatments for this devastating disease. Second, while MAC accumulation was clearly associated with heightened oxidative-stress induced cytotoxicity in high-risk CECs, the precise mechanisms, whether caused by compromised CFH binding or intracellular complement dysfunction, require further elucidation.

In summary, this study provides evidence that the high-risk CFH 402H variant sensitises choriocapillaris endothelial cells to environmental stressors, leading to an increase in oxidative stress and triggering complement-mediated injury and cell death. This is associated with extensive stress-induced reprogramming, including dysregulation of lipid metabolism in both high-risk CECs, complement activation, angiogenic signalling, and endothelial-to-mesenchymal transition pathways imbalance, highly pronounced in HR_1. These findings align with published studies providing analyses of the AMD eyes showing diminished choriocapillaris network in the early stage of the disease, coupled with pathological neovascularisation in wet AMD, highlighting the choriocapillaris as a critical and overlooked therapeutic target in preventing AMD initiation and progression.

## Materials and methods

### Generation of isogenic controls for high-risk (*CFH*: Y402H) iPSC lines

Both high-risk iPSC (Y402H) lines, generated from dermal fibroblasts of wet AMD patients as described in (Hallam et al., 2017), were electroporated using a 4D-Nucleofector (Lonza) machine to transfer gRNA/Cas9 and ssODN directly into the nucleus. gRNA (AATGGATATAATCAAAATCA) and ssODN were designed with Benchling tools and synthesised by Thermo Fisher Scientific and Integrated DNA Technologies, respectively. TrueCut Cas9 Protein V2 (Thermo Fisher Scientific, A36499) was incubated with gRNA for 10 minutes, followed by the addition of ssODN in 100 µl nucleofection solution. Cells were grown for the last 24 hours before nucleofection in the presence of a nonhomologous end joining (NHEJ) inhibitor (SCR7 pyrazine, Sigma, SML1546, at a concentration of 10 μM) to stimulate homology-directed repair (HDR). iPSCs were dissociated for nucleofection with StemPro® Accutase® (Gibco, A11105-01). The ribonucleoprotein complexes alongside ssODN were delivered to the cells using the CB150 program at a 1:2 ratio of Cas9 Nuclease to gRNA/ssODN. After electroporation, cells were mixed with 500 µl mTeSR1 medium (Stem Cell Technologies, 85850) containing 10 μM ROCK inhibitor (Y27632, Tocris) and SCR7 pyrazine (10 μM). The cell suspension was distributed among at least five Matrigel (Corning, 354230) coated 10 cm plates supplemented with 4 ml of TeSR1 medium, 10 μM ROCK inhibitor and SCR7 pyrazine (10 μM). Medium was replaced 48 hours post-seeding with mTeSR1 only and changed every second day afterwards. Colonies were picked manually around 10 days later and grown in 24-well plates to be assessed for gene editing with restriction enzyme assay (*FatI*, NEB, R0650L). Selected clones were sequenced to confirm correction of high-risk C nucleotide to T (H402Y) in the *CFH* gene (F: GCACATAAGTGATTACACCTGTC, R: TGTAACTGTGGTCTGCGCTT). Primers used to obtain PCR products enclosing off-target sites, with the lowest number of mismatches, identified with Off-Spotter, were sequenced to exclude additional changes to the genome (**Table S1)**.

### Generation of iPSC-CECs

The differentiation plan followed the (Mulfaul et al., 2020) published protocol. Briefly, upon reaching 80% confluency (day 0) iPSC colonies were lifted off gently with Versene, EDTA (Life Technologies, 15040066) from 3-wells of Matrigel (Corning, 354230) coated 6-well plate (TPP), resuspend in a mix of 6 ml mTeSR1 and 3 ml Endothelial Cell Growth Media (R&D Systems, CCM027) with 10 μM ROCK inhibitor (Y27632, Tocris) and transferred to 2-wells of ultra-low attachment 6-well plate (Corning, 3471). The following day (day 1), embryonic bodies (EBs) were harvested and resuspended in a mix of 3 ml mTeSR1 and 6 ml Endothelial Cell Growth Media supplemented with 20 ng/ml BMP-4 (Life Technologies, PHC9531). On day 2, medium was changed to 9 ml of Endothelial Cell Growth Media with 20 ng/ml BMP-4 and 10 ng/ml activin A (Biotechne, 338-AC-010). Medium for day 3 was supplemented with 20 ng/ml BMP-4, 10 ng/ml activin A and 8 ng/ml FGF-2 (Life Technologies, GMP100-18B-100UG).

On day 5 EBs were distributed over 3 wells of Matrigel (Corning, 354230) coated 6-well plate in Endothelial Cell Growth Media with 10 ng/ml BMP4, 8 ng/ml FGF-2, 25 ng/ml VEGF (Life Technologies, 100-20) and 25 ng/ml CTGF (Life Technologies, 120-19), followed by the same media changed on day 6. Up to day 14, Endothelial Cell Growth Media with 8 ng/ml FGF-2, 25 ng/ml VEGF and 25 ng/ml CTGF were changed every second day. On day 14, after cell treatment with TrypLe Express (Life Technologies, 12604013), CD31-positive cells were isolated from a single cell suspension with the CD31 MicroBead Kit (Miltenyi Biotec, 130-091-935) according to the manufacturer’s protocol. In summary, CD31 MicroBeads immunolabeled cells in PBS containing 0.04% non-acetylated BSA (Scientific Laboratory Supplies, B6917-25MG) were separated with an LS column (Miltenyi Biotec, 130-042-401) placed in the magnetic field of a MACS Separator. Magnetically captured CD31^+^ cells were eluted with Endothelial Cell Growth Media supplemented with 8 ng/ml FGF-2, 25 ng/ml VEGF and 25 ng/ml CTGF and cultured on Matrigel-coated 6-well plates.

### Cell maintenance

Both iPSCs and CECs were cultured in Matrigel-coated plates and mTeSR1 growth media or Endothelial Cell Growth Media supplemented with 8 ng/ml FGF-2, 25 ng/ml VEGF and 25 ng/ml CTGF with 1% Pen/Strep (Life Technologies, 15140-122) in a humidified, 5% CO_2_, 37°C incubator. iPSCs were passaged in a ratio of 1:6 twice per week using Versene (EDTA), whereas CECs were passaged in a ratio of 1:3 when reaching around 100% confluency with TrypLe Express.

### Western blotting

Supernatants were collected from CEC cultures in 12-well plates after 4 days of incubation with Endothelial Cell Growth Media supplemented with 8 ng/ml FGF-2, 25 ng/ml VEGF and 25 ng/ml CTGF, then centrifuged at 4°C in an Eppendorf centrifuge 5415 to remove cell debris. The supernatants were subsequently concentrated with Amicon Ultra-0.5ml centrifugal filters (Merck, UFC500396) according to the manufacturer’s protocol.

CECs to assess autophagy flux were incubated for 24 hours with 500nM rapamycin (Invivogen, tlrl-rap-5) and 100nM bafilomycin A1 (Invivogen, tlrl-baf1) over the last 4 hours. Cell lysates were prepared from pelleted CECs by the addition of cold RIPA Lysis Buffer (MILLIPORE, 20-188) with protease inhibitor mix (EDTA-free, Roche), followed by 30-minute incubation on ice and pipetting in 15-minute intervals. Supernatants were retained after 5-minute centrifugation (4°C) for SDS-PAGE. Between 10 to 20 µg of proteins from the cell lysates or 50 µg for supernatants were mixed with NuPAGE LDS Sample Buffer and NuPAGE Sample Reducing Agent (Thermofisher Scientific, NP0004), which was omitted in the case of CFI protein detection. Samples were incubated at 70°C for 10 minutes and separated on NuPage 4-12% Bis Tris Gel (Cell Signalling Technology, 125395) in Novex NuPAGE MOPS SDS Running Buffer alongside PageRuler™ Plus Prestained Protein Ladder (Thermo Scientific, 26619). The proteins were transferred to PVDF membranes by iBlot2 Life Technologies Apparatus. The primary antibodies (**Table S2**) were incubated with the membranes overnight, at 4°C, followed by secondary horseradish peroxidase-conjugated anti-rabbit, anti-mouse IgG (Dako), or anti-sheep IgG H&L (Abcam) at room temperature for 1 hour. The signal was obtained using the Thermo Fisher Scientific Super Signal West Pico Plus Chemiluminescent substrate kit and detected by Amersham Imager 600. Uncropped original Western blot images are provided in the supplementary materials.

### Immunohistochemistry

The CECs were plated on Matrigel-coated 4-well glass chamber slides (Thermo Scientific, 154526) and maintained with medium changes every second day. Once at the required confluency, the medium was removed, and cells were briefly washed with PBS, then fixed in 4% paraformaldehyde for 10 minutes. Three subsequent washes with PBS were performed after removal of fixative. CECs were blocked and permeabilised for 1 hour, at room temperature (RT) in 0.05% Triton-X-100 (Sigma) and 5% Normal Goat Serum (Thermo Scientific) and 3% bovine serum albumin (Sigma, A2153-100G) in PBS. After removal of the blocking solution, the primary antibody was applied at concentrations shown in **Table S2** in blocking buffer overnight at 4°C. Cells were washed three times with PBS after overnight incubation and incubated with secondary antibodies: donkey anti-mouse 546 (Life Technologies, A10036) or donkey anti-rabbit 546 (Invitrogen, A10040) in PBS for 2 hours at RT. Cells were washed again and mounted in Vecta-shield (Vector Labs, Burlingame, CA) with Hoechst 33342 (1:1000, Thermo Scientific). Fluorescence microscopy images of CECs were taken using a Zeiss Axio Imager Z2 equipped with an Apotome2 (Zeiss, Germany). Secondary antibody controls were performed in the same manner, but with the primary antibody addition step omitted. Quantitative analyses of immunoassayed sections were carried out using ZEN (blue edition; ZEISS) and ImageJ.

### ELISA

To measure the total amount of C3b, iC3b, and C3c in tissue culture supernatant, an ELISA was established that captured these fragments using mAb bH6 (Hycult Biotech, The Netherlands; 2 μg/mL coat); C3b (Complement Technology, Tyler, Texas) was used for the standard curve. Plates were blocked with phosphate-buffered saline (PBS), 5% nonfat milk (wt/vol), 0.1% Tween 20. Samples were diluted in PBS, 1% bovine serum albumin (BSA) (wt/vol), 10 m mM EDTA. Bound C3 fragments were detected using a polyclonal rabbit anti-human C3 (DAKO, A0063) at 1:1000 dilution, followed by goat anti-rabbit Ig-HRPO (Invitrogen, 65-6120) 1:1000. The assay was developed using TMB and absorbance was measured at 450 nm. Values were interpolated from the standard curve.

To measure total uncleaved C3, an ELISA was established that captured the C3a domain using mAb Clone2898 (HM2075, Hycult Biotech; 2 μg/mL coat). Complement C3 (Complement Technologies, A113c) was used for the standard curve. Plates were blocked with PBS/10 mM EDTA, 3% BSA (wt/vol). Samples were diluted in PBS, 1% BSA (wt/vol), 10 m mM EDTA. Bound C3 was detected using a polyclonal goat anti-C3 (Complement Technology, A213) at 1:5000 dilution, followed by donkey anti-goat Ig-HRPO (Jackson Immuno, 705-035-147) 1:8000. The assay was developed using TMB and absorbance was measured at 450 nm. Values were interpolated from the standard curve.

### Transmission Electron Microscopy

The transwell inserts (Greiner Bio-One, 0.4 µm pore, translucent, 24-well, 662641) with CECs were briefly washed with PBS and fixed with 2% glutaraldehyde at 4°C. Cell processing and imaging were performed at Newcastle University Electron Microscopy Research Services. Ultrathin sections were stained with heavy metal salts (uranyl acetate and lead citrate) and imaged on a Hitachi HT7800 120kV TEM using an EMSIS CMOS Xarosa high-resolution camera (Hitachi, Japan).

### Quantitative RT-PCR

RNA was extracted from frozen cell pellets using the ReliaPrep RNA Cell Miniprep System (Promega) according to the manufacturer’s instructions. cDNA synthesis was performed using the GoScript Reverse Transcription System (Promega). qPCRs were performed with a QuantStudio 7 Flex Real-Time PCR System (Applied Biosystems) using SYBR green reaction technology (Promega). Ct results of the target genes were normalised to the Ct of the reference gene GAPDH (ΔCt); the ΔCt obtained was then normalised to the ΔCt of the control sample (ISO), generating the ΔΔCt, allowing determination of fold difference (2^-ΔΔCt^). A list of the primers used is shown in **Table S3**.

### Cell barrier function formation

Barrier formation was measured using the xCELLigence system (Agilent). Cells were plated in a 16-well plate (Agilent) at a 2×10^4^ cells per well in 200 μl media (R&D). Cells were left to settle in the plate for 15 minutes at RT in the safety cabinet before starting barrier measurements. Barrier capacity was measured as cell index; a higher cell index indicates higher barrier capacity. The instrument was set to record one cell index measurement every 15 minutes. Cells were left to form a barrier overnight or until a plateau was reached. At that point, the experiment was interrupted for analysis.

### Cellular respiration

Glycolysis and mitochondrial respiration were measured using the Seahorse XFe96 analyser (Agilent, UK). Cells were seeded in an XFe96 Pro cell culture microplate at 20,000 cells per well and allowed to form a monolayer overnight. The XFe96 Pro sensor cartridges were hydrated overnight according to the manufacturer’s instructions. On the day of the experiment, the cell culture media were replaced with Seahorse assay media (103575–100, Agilent). Glycolysis was tested using Agilent Seahorse Cell Glycolysis Stress Test Kit (103020-100, Agilent) and Mitochondrial respiration was assessed using the Agilent Seahorse Cell Mito Stress Test kit (103015-100, Agilent). Data was analysed using Seahorse Wave Controller Software 2.6.1 and GraphPad Prism.

### Endothelial Cell Tube Formation Assay *in vitro*

#### Tubulogenesis assay

This was performed using μ-slides 15 Well 3D ibiTreat (81506). Matrigel was mixed with media (R&D) in a ratio of 60:40 on ice and used to resuspend cells. 1.5×10^4^ cells were plated in a volume of 10 µl per well. Slides were left for 40 min in the incubator to allow polymerisation of Matrigel. Subsequently, wells were filled with 50 μL of media and left to form tubes for 24 hours. Cells were stained with Calcein for 1h at 37 °C and washed twice with media before imaging. Entire wells were imaged using the EVOS imaging system, and composite images were created.

To investigate the effect of Hydroquinone on HR and their ISO CECs, 5 × 10⁴ cells were resuspended in Corning Matrigel (356231) at a 60:40 ratio and dispensed as 50 µL droplets into five wells per condition of a 96 well plate. The Matrigel–cell droplets were allowed to polymerize at 37 °C for 60 min, after which appropriate prewarmed culture medium was gently added to each well, and the plate was incubated overnight under standard culture conditions. The HQ (200 µM) was added after 24 hours to the respective wells. Cells were subsequently stained with Calcein for 30 minutes at 37 °C, and washed with culture medium, and the entire well area was imaged using the Zeiss Apotome imaging system. Images were further analysed in FIJI using the Vessel Analysis package to quantify vessel density and vessel length density, while vessel thickness was quantified manually from the same regions of interest.

#### Cytotoxicity assays

To assess the cytotoxicity effect of human serum (MERCK, H4522-20ml), high-risk RPE conditioned medium, and Hydroquinone (Merck, H9003), CECs were seeded in Matrigel-coated 96-well plates (PPT). Media were changed every other day during 96 hr incubation.

For the Incucyte® Cytotoxicity Assay, the Incucyte® Cytotox NIR Dye (Sartorius, 4846) was used, allowing real-time measurements of cell death based on cell membrane integrity. Briefly, CECs were monitored in Incucyte (Sartorius, IC70795) in the presence of Cytotox NIR Dye at a dilution of 1:1000. The number of dead, fluorescently labelled cells was detected and quantified by Incucyte image analysis software.

#### Determination of ROS levels

Levels of reactive oxygen species (ROS) were measured using ROS-Glo™ H_2_O_2_ Assay (Promega, G8820) according to the manufacturer’s protocol. Briefly, CECs were incubated with the 200 µM Hydroquinone and hydrogen peroxide substrate for 3 hr at 37°C, followed by the addition of ROS-Glo™ Detection Solution and an incubation for 20 minutes at RT. Luminescent signal was measured by a GloMax Discover plate reader.

#### RNA-Seq analysis

RNA integrity from 24 samples was assessed using Agilent Broad Range ScreenTape assays on the Agilent 4200 TapeStation. All samples demonstrated high RNA quality, with RNA integrity number (RIN) values ≥7. mRNA sequencing libraries were prepared using the Illumina Stranded mRNA Library Prep protocol, in which poly(dT) beads were used to enrich polyadenylated mRNA transcripts. Libraries were subsequently evaluated using D1000 ScreenTape assays on the Agilent 4200 TapeStation and quantified using the Qubit dsDNA Broad Range Assay Kit. All 24 libraries were pooled in equimolar concentrations and sequenced on the Illumina NovaSeq 6000 using an S1 flow cell with the 200-cycle v1.5 reagent kit. Sequencing was performed with paired-end 101 bp reads and dual 10 bp index reads (101/10/10/101 cycles), with 5% PhiX control library spike-in.

Sequencing reads (FASTQ files) for all samples (n=24) were quality controlled, aligned and quantified using the nf-core/rnaseq pipeline (v.3.22.2) and the GRCh38 reference genome (Ensemble Release 115). nf-core/rnaseq was run using the default “STAR Salmon” workflow (Harshil Patel et al., 2026). Transcript count matrices were imported into the R statistical computing environment ((R Core Team 2025), v.4.5.2) for analysis with DESeq2 ((Love et al., 2014), v.1.48.2). After normalisation with DESeq2, suitable contrasts were selected for analysis, following best practice workflows (https://github.com/bcbio/rnaseq-reports). Volcano plots of DEG were produced using The EnhancedVolcano R package (v1.26.0) and Gene Set Enrichment Analyses (GSEA) were completed using the R packages clusterProfiler (v.4.16.0) and fgsea (v.1.34.2), and the DAVID web server (Sherman et al., 2022).

Raw RNA-Seq data have been deposited within the Gene Expression Omnibus (**GEO**) repository and can be accessed using the reference ID: GSE333292

## Statistical analysis

An unpaired two-tailed Student’s t-test or one/two-way ANOVA was used to compare the mean ± SEM values between high-risk and ISO CECs. The analyses were performed with GraphPad Prism software; values of p ≤ 0.05 were considered statistically significant (*p ≤0.05, **p ≤ 0.01, ***p ≤ 0.001, ****p ≤ 0.0001).

## Acknowledgements

The authors are grateful for financial support from the Macular Society, EPSRC/ERC (EP/Y031016/1), Leverhulme Trust (RPG-2023-149) and BBSRC UK/Japan (BB/W018470/2) for funding this work. The purchase of IncuCyte used in this study was supported by a UKRI MRC Capital Funding for World Class Labs Award (MR/X012360/1).

## Competing interests

The authors declare no conflict of interest.

## Author contributions

**AR**: data acquisition, data analysis, figure preparation, manuscript writing and dissemination

**JDF, VP, PMB, HD, DB:** data acquisition and analysis

**MT**: bioinformatic analysis, contributed to figures preparation and manuscript writing

**JK, RQ, JC**: data acquisition

**KM, DK, MKA**: study design and provided reagents for this work

**LA, RM**: study design and fund raising

**ML**: study design, manuscript writing and fund raising, dissemination. All authors read and approved the final manuscript.

## Notes

### Competing Interest Statement

The authors have declared no competing interest.

## References

Armento, A., Honisch, S., Panagiotakopoulou, V., Sonntag, I., Jacob, A., Bolz, S., Kilger, E., Deleidi, M., Clark, S., Ueffing, M., 2020. Loss of Complement Factor H impairs antioxidant capacity and energy metabolism of human RPE cells. Sci. Rep. 10, 10320. 10.1038/s41598-020-67292-z

Armento, A., Sonntag, I., Almansa-Garcia, A.-C., Sen, M., Bolz, S., Arango-Gonzalez, B., Kilger, E., Sharma, R., Bharti, K., Fernandez-Godino, R., de la Cerda, B., Clark, S.J., Ueffing, M., 2025. The AMD-associated genetic polymorphism CFH Y402H confers vulnerability to Hydroquinone-induced stress in iPSC-RPE cells. Front. Immunol. 16, 1527018. 10.3389/fimmu.2025.1527018

Armento, A., Ueffing, M., Clark, S.J., 2021. The complement system in age-related macular degeneration. Cell. Mol. Life Sci. CMLS 78, 4487–4505. 10.1007/s00018-021-03796-9

Balaiya, S., Khetpal, V., Chalam, K.V., 2012. Hypoxia initiates sirtuin1-mediated vascular endothelial growth factor activation in choroidal endothelial cells through hypoxia inducible factor-2α. Mol. Vis. 18, 114–120.

Berenberg, T.L., Metelitsina, T.I., Madow, B., Dai, Y., Ying, G.-S., Dupont, J.C., Grunwald, L., Brucker, A.J., Grunwald, J.E., 2012. The association between drusen extent and foveolar choroidal blood flow in age-related macular degeneration. Retina Phila. Pa 32, 25–31. 10.1097/IAE.0b013e3182150483

Blatt, A.Z., Pathan, S., Ferreira, V.P., 2016. Properdin: a tightly regulated critical inflammatory modulator. Immunol. Rev. 274, 172–190. 10.1111/imr.12466

Cabral, T., Mello, L.G.M., Lima, L.H., Polido, J., Regatieri, C.V., Belfort, R., Mahajan, V.B., 2017. Retinal and choroidal angiogenesis: a review of new targets. Int. J. Retina Vitr. 3, 31. 10.1186/s40942-017-0084-9

Cáceres-Vélez, P.R., Hui, F., Hercus, J., Bui, B., Jusuf, P.R., 2022. Restoring the oxidative balance in age-related diseases - An approach in glaucoma. Ageing Res. Rev. 75, 101572. 10.1016/j.arr.2022.101572

Campagna, R., Mazzanti, L., Pompei, V., Alia, S., Vignini, A., Emanuelli, M., 2024. The Multifaceted Role of Endothelial Sirt1 in Vascular Aging: An Update. Cells 13, 1469. 10.3390/cells13171469

Castilho, Á.F., Aveleira, C.A., Leal, E.C., Simões, N.F., Fernandes, C.R., Meirinhos, R.I., Baptista, F.I., Ambrósio, A.F., 2012. Heme oxygenase-1 protects retinal endothelial cells against high glucose- and oxidative/nitrosative stress-induced toxicity. PloS One 7, e42428. 10.1371/journal.pone.0042428

Cerniauskas, E., Kurzawa-Akanbi, M., Xie, L., Hallam, D., Moya-Molina, M., White, K., Steel, D., Doherty, M., Whitfield, P., Al-Aama, J., Armstrong, L., Kavanagh, D., Lambris, J.D., Korolchuk, V.I., Harris, C., Lako, M., 2020. Complement modulation reverses pathology in Y402H-retinal pigment epithelium cell model of age-related macular degeneration by restoring lysosomal function. Stem Cells Transl. Med. 9, 1585–1603. 10.1002/sctm.20-0211

Chen, X., Xu, Y., Ju, Y., Gu, P., 2024. Metabolic Regulation of Endothelial Cells: A New Era for Treating Wet Age-Related Macular Degeneration. Int. J. Mol. Sci. 25, 5926. 10.3390/ijms25115926

Chirco, K.R., Sohn, E.H., Stone, E.M., Tucker, B.A., Mullins, R.F., 2017. Structural and molecular changes in the aging choroid: implications for age-related macular degeneration. Eye 31, 10–25. 10.1038/eye.2016.216

Curcio, C.A., Johnson, M., Rudolf, M., Huang, J.-D., 2011. The oil spill in ageing Bruch membrane. Br. J. Ophthalmol. 95, 1638–1645. 10.1136/bjophthalmol-2011-300344

de Córdoba, S.R., de Jorge, E.G., 2008. Translational mini-review series on complement factor H: genetics and disease associations of human complement factor H. Clin. Exp. Immunol. 151, 1–13. 10.1111/j.1365-2249.2007.03552.x

Ebeling, M.C., Geng, Z., Kapphahn, R.J., Roehrich, H., Montezuma, S.R., Dutton, J.R., Ferrington, D.A., 2021. Impaired Mitochondrial Function in iPSC-Retinal Pigment Epithelium with the Complement Factor H Polymorphism for Age-Related Macular Degeneration. Cells 10, 789. 10.3390/cells10040789

Eklund, L., Piuhola, J., Komulainen, J., Sormunen, R., Ongvarrasopone, C., Fássler, R., Muona, A., Ilves, M., Ruskoaho, H., Takala, T.E., Pihlajaniemi, T., 2001. Lack of type XV collagen causes a skeletal myopathy and cardiovascular defects in mice. Proc. Natl. Acad. Sci. U. S. A. 98, 1194–1199. 10.1073/pnas.98.3.1194

Epstein, T., Xu, L., Gillies, R.J., Gatenby, R.A., 2014. Separation of metabolic supply and demand: aerobic glycolysis as a normal physiological response to fluctuating energetic demands in the membrane. Cancer Metab. 2, 7. 10.1186/2049-3002-2-7

Febbraio, M., Hajjar, D.P., Silverstein, R.L., 2001. CD36: a class B scavenger receptor involved in angiogenesis, atherosclerosis, inflammation, and lipid metabolism. J. Clin. Invest. 108, 785–791. 10.1172/JCI14006

Ferrington, D.A., Ebeling, M.C., Kapphahn, R.J., Terluk, M.R., Fisher, C.R., Polanco, J.R., Roehrich, H., Leary, M.M., Geng, Z., Dutton, J.R., Montezuma, S.R., 2017. Altered bioenergetics and enhanced resistance to oxidative stress in human retinal pigment epithelial cells from donors with age-related macular degeneration. Redox Biol. 13, 255–265. 10.1016/j.redox.2017.05.015

Ferrington, D.A., Fisher, C.R., Kowluru, R.A., 2020. Mitochondrial Defects Drive Degenerative Retinal Diseases. Trends Mol. Med. 26, 105–118. 10.1016/j.molmed.2019.10.008

Fisher, C.R., Ferrington, D.A., 2018. Perspective on AMD Pathobiology: A Bioenergetic Crisis in the RPE. Investig. Opthalmology Vis. Sci. 59, AMD41. 10.1167/iovs.18-24289

Fritsche, L.G., Fariss, R.N., Stambolian, D., Abecasis, G.R., Curcio, C.A., Swaroop, A., 2014. Age-related macular degeneration: genetics and biology coming together. Annu. Rev. Genomics Hum. Genet. 15, 151–171. 10.1146/annurev-genom-090413-025610

Fu, Z., Kern, T.S., Hellström, A., Smith, L.E.H., 2021. Fatty acid oxidation and photoreceptor metabolic needs. J. Lipid Res. 62, 100035. 10.1194/jlr.TR120000618

GBD 2021 Global AMD Collaborators, 2025. Global burden of vision impairment due to age-related macular degeneration, 1990-2021, with forecasts to 2050: a systematic analysis for the Global Burden of Disease Study 2021. Lancet Glob. Health 13, e1175–e1190. 10.1016/S2214-109X(25)00143-3

Go, Y.-M., Jones, D.P., 2008. Redox compartmentalization in eukaryotic cells. Biochim. Biophys. Acta 1780, 1273–1290. 10.1016/j.bbagen.2008.01.011

Go, Y.-M., Zhang, J., Fernandes, J., Litwin, C., Chen, R., Wensel, T.G., Jones, D.P., Cai, J., Chen, Y., 2020. MTOR-initiated metabolic switch and degeneration in the retinal pigment epithelium. FASEB J. Off. Publ. Fed. Am. Soc. Exp. Biol. 34, 12502–12520. 10.1096/fj.202000612R

Grebe, R., Mughal, I., Bryden, W., McLeod, S., Edwards, M., Hageman, G.S., Lutty, G., 2019. Ultrastructural analysis of submacular choriocapillaris and its transport systems in AMD and aged control eyes. Exp. Eye Res. 181, 252–262. 10.1016/j.exer.2019.02.018

Hallam, D., Collin, J., Bojic, S., Chichagova, V., Buskin, A., Xu, Y., Lafage, L., Otten, E.G., Anyfantis, G., Mellough, C., Przyborski, S., Alharthi, S., Korolchuk, V., Lotery, A., Saretzki, G., McKibbin, M., Armstrong, L., Steel, D., Kavanagh, D., Lako, M., 2017. An Induced Pluripotent Stem Cell Patient Specific Model of Complement Factor H (Y402H) Polymorphism Displays Characteristic Features of Age-Related Macular Degeneration and Indicates a Beneficial Role for UV Light Exposure. Stem Cells Dayt. Ohio 35, 2305–2320. 10.1002/stem.2708

Harshil Patel, Jonathan Manning, Phil Ewels, Maxime U Garcia, Alexander Peltzer, Rickard Hammarén, Olga Botvinnik, Adam Talbot, Friederike Hanssen, Gregor Sturm, nf-core bot, Matthias Zepper, Denis Moreno, Ezra Greenberg, Pranathi Vemuri, Gary Burnett, Mahesh Binzer-Panchal, silviamorins, Lorena Pantano, Robert Syme, EladH1, Gavin Kelly, Lorenzo Sola, James A. Fellows Yates, Matthias Hörtenhuber, Gabriel Lichtenstein, Edmund Miller, Jose Espinosa-Carrasco, rfenouil, Luke Zappia, 2026. nf-core/rnaseq: nf-core/rnaseq v3.23.0 - Gallium Gecko. 10.5281/ZENODO.1400710

He, J., Ma, M., Li, D., Wang, K., Wang, Q., Li, Qiuguo, He, H., Zhou, Y., Li, Qinglong, Hou, X., Yang, L., 2021. Sulfiredoxin-1 attenuates injury and inflammation in acute pancreatitis through the ROS/ER stress/Cathepsin B axis. Cell Death Dis. 12, 626. 10.1038/s41419-021-03923-1

Henning, Y., Willbrand, K., Larafa, S., Weißenberg, G., Matschke, V., Theiss, C., Görtz, G.-E., Matschke, J., 2023. Cigarette smoke causes a bioenergetic crisis in RPE cells involving the downregulation of HIF-1α under normoxia. Cell Death Discov. 9, 398. 10.1038/s41420-023-01695-5

Hu, M., Ladowski, J.M., Xu, H., 2024. The Role of Autophagy in Vascular Endothelial Cell Health and Physiology. Cells 13, 825. 10.3390/cells13100825

Huang, J., Gu, S., Chen, M., Zhang, S.-J., Jiang, Z., Chen, X., Jiang, C., Liu, G., Radu, R.A., Sun, X., Vollrath, D., Du, J., Yan, B., Zhao, C., 2019. Abnormal mTORC1 signaling leads to retinal pigment epithelium degeneration. Theranostics 9, 1170–1180. 10.7150/thno.26281

Jakobsen, T.S., Adsersen, R.L., Askou, A.L., Corydon, T.J., 2024. Functional Roles of Pigment Epithelium-Derived Factor in Retinal Degenerative and Vascular Disorders: A Scoping Review. Invest. Ophthalmol. Vis. Sci. 65, 41. 10.1167/iovs.65.14.41

Jiang, F., Ma, J., Lei, C., Zhang, Y., Zhang, M., 2025. Age-Related Macular Degeneration: Cellular and Molecular Signaling Mechanisms. Int. J. Mol. Sci. 26, 6174. 10.3390/ijms26136174

Johnson, P.T., Betts, K.E., Radeke, M.J., Hageman, G.S., Anderson, D.H., Johnson, L.V., 2006. Individuals homozygous for the age-related macular degeneration risk-conferring variant of complement factor H have elevated levels of CRP in the choroid. Proc. Natl. Acad. Sci. U. S. A. 103, 17456–17461. 10.1073/pnas.0606234103

Kouser, L., Abdul-Aziz, M., Nayak, A., Stover, C.M., Sim, R.B., Kishore, U., 2013. Properdin and factor h: opposing players on the alternative complement pathway “see-saw.” Front. Immunol. 4, 93. 10.3389/fimmu.2013.00093

Kumari, R., Deshmukh, R.S., Das, S., 2019. Caspase-10 inhibits ATP-citrate lyase-mediated metabolic and epigenetic reprogramming to suppress tumorigenesis. Nat. Commun. 10, 4255. 10.1038/s41467-019-12194-6

Kurzawa-Akanbi, M., Whitfield, P., Burté, F., Bertelli, P.M., Pathak, V., Doherty, M., Hilgen, B., Gliaudelytė, L., Platt, M., Queen, R., Coxhead, J., Porter, A., Öberg, M., Fabrikova, D., Davey, T., Beh, C.S., Georgiou, M., Collin, J., Boczonadi, V., Härtlova, A., Taggart, M., Al-Aama, J., Korolchuk, V.I., Morris, C.M., Guduric-Fuchs, J., Steel, D.H., Medina, R.J., Armstrong, L., Lako, M., 2022. Retinal pigment epithelium extracellular vesicles are potent inducers of age-related macular degeneration disease phenotype in the outer retina. J. Extracell. Vesicles 11, 12295. 10.1002/jev2.12295

Lee, D., Xu, I.M.-J., Chiu, D.K.-C., Leibold, J., Tse, A.P.-W., Bao, M.H.-R., Yuen, V.W.-H., Chan, C.Y.-K., Lai, R.K.-H., Chin, D.W.-C., Chan, D.F.-F., Cheung, T.-T., Chok, S.-H., Wong, C.-M., Lowe, S.W., Ng, I.O.-L., Wong, C.C.-L., 2019. Induction of Oxidative Stress Through Inhibition of Thioredoxin Reductase 1 Is an Effective Therapeutic Approach for Hepatocellular Carcinoma. Hepatology 69, 1768–1786. 10.1002/hep.30467

Lejoyeux, R., Benillouche, J., Ong, J., Errera, M.-H., Rossi, E.A., Singh, S.R., Dansingani, K.K., da Silva, S., Sinha, D., Sahel, J.-A., Freund, K.B., Sadda, S.R., Lutty, G.A., Chhablani, J., 2022. Choriocapillaris: Fundamentals and advancements. Prog. Retin. Eye Res. 87, 100997. 10.1016/j.preteyeres.2021.100997

Li, H., Liu, B., Lian, L., Zhou, J., Xiang, S., Zhai, Y., Chen, Y., Ma, X., Wu, W., Hou, L., 2021. High dose expression of heme oxigenase-1 induces retinal degeneration through ER stress-related DDIT3. Mol. Neurodegener. 16, 16. 10.1186/s13024-021-00437-4

Li, M., Wang, P., Huo, S.T., Qiu, H., Li, C., Lin, S., Guo, L., Ji, Y., Zhu, Y., Liu, J., Guo, J., Na, J., Hu, Y., 2024. Human Pluripotent Stem Cells Derived Endothelial Cells Repair Choroidal Ischemia. Adv. Sci. Weinh. Baden-Wurtt. Ger. 11, e2302940. 10.1002/advs.202302940

Li, S., Yuan, H., Li, L., Li, Q., Lin, P., Li, K., 2025. Oxidative Stress and Reprogramming of Lipid Metabolism in Cancers. Antioxidants 14, 201. 10.3390/antiox14020201

Lin, Y., Gahn, J., Banerjee, K., Dobreva, G., Singhal, M., Dubrac, A., Ola, R., 2024. Role of endothelial PDGFB in arterio-venous malformations pathogenesis. Angiogenesis 27, 193–209. 10.1007/s10456-023-09900-w

Liu, K., Chen, L.J., Lai, T.Y.Y., Tam, P.O.S., Ho, M., Chiang, S.W.Y., Liu, D.T.L., Young, A.L., Yang, Z., Pang, C.P., 2014. Genes in the high-density lipoprotein metabolic pathway in age-related macular degeneration and polypoidal choroidal vasculopathy. Ophthalmology 121, 911–916. 10.1016/j.ophtha.2013.10.042

Love, M.I., Huber, W., Anders, S., 2014. Moderated estimation of fold change and dispersion for RNA-seq data with DESeq2. Genome Biol. 15, 550. 10.1186/s13059-014-0550-8

Maloney, S.C., Antecka, E., Granner, T., Fernandes, B., Lim, L.-A., Orellana, M.E., Burnier, M.N., 2013. Expression of SIRT1 in choroidal neovascular membranes. Retina 33, 862–866. 10.1097/IAE.0b013e31826af556

Mameli, E., Martello, A., Caporali, A., 2022. Autophagy at the interface of endothelial cell homeostasis and vascular disease. FEBS J. 289, 2976–2991. 10.1111/febs.15873

Margolis, R., Spaide, R.F., 2009. A pilot study of enhanced depth imaging optical coherence tomography of the choroid in normal eyes. Am. J. Ophthalmol. 147, 811–815. 10.1016/j.ajo.2008.12.008

McAleese, C., Joudah, G., Salt, I.P., Petrie, J.R., Leiper, J.M., Dowsett, L.B., 2025. Heterogeneous metabolic response of endothelial cells from different vascular beds to experimental hyperglycaemia and metformin. J. Physiol. 10.1113/JP288006

Mitchell, P., Liew, G., Gopinath, B., Wong, T.Y., 2018. Age-related macular degeneration. Lancet Lond. Engl. 392, 1147–1159. 10.1016/S0140-6736(18)31550-2

Mitter, S.K., Song, C., Qi, X., Mao, H., Rao, H., Akin, D., Lewin, A., Grant, M., Dunn, W., Ding, J., Bowes Rickman, C., Boulton, M., 2014. Dysregulated autophagy in the RPE is associated with increased susceptibility to oxidative stress and AMD. Autophagy 10, 1989–2005. 10.4161/auto.36184

Mori, K., Ishikawa, K., Fukuda, Y., Ji, R., Wada, I., Kubo, Y., Akiyama, M., Notomi, S., Murakami, Y., Nakao, S., Arakawa, S., Shiose, S., Hisatomi, T., Yoshida, S., Kannan, R., Sonoda, K.-H., 2022. TNFRSF10A downregulation induces retinal pigment epithelium degeneration during the pathogenesis of age-related macular degeneration and central serous chorioretinopathy. Hum. Mol. Genet. 31, 2194–2206. 10.1093/hmg/ddac020

Mrejen, S., Spaide, R.F., 2013. Optical coherence tomography: imaging of the choroid and beyond. Surv. Ophthalmol. 58, 387–429. 10.1016/j.survophthal.2012.12.001

Mulfaul, K., Giacalone, J.C., Voigt, A.P., Riker, M.J., Ochoa, D., Han, I.C., Stone, E.M., Mullins, R.F., Tucker, B.A., 2020. Stepwise differentiation and functional characterization of human induced pluripotent stem cell-derived choroidal endothelial cells. Stem Cell Res. Ther. 11, 409. 10.1186/s13287-020-01903-4

Mulfaul, K., Russell, J.F., Voigt, A.P., Stone, E.M., Tucker, B.A., Mullins, R.F., 2022. The Essential Role of the Choriocapillaris in Vision: Novel Insights from Imaging and Molecular Biology. Annu. Rev. Vis. Sci. 8, 33–52. 10.1146/annurev-vision-100820-085958

Mullins, R.F., Dewald, A.D., Streb, L.M., Wang, K., Kuehn, M.H., Stone, E.M., 2011a. Elevated membrane attack complex in human choroid with high risk complement factor H genotypes. Exp. Eye Res. 93, 565–567. 10.1016/j.exer.2011.06.015

Mullins, R.F., Johnson, M.N., Faidley, E.A., Skeie, J.M., Huang, J., 2011b. Choriocapillaris vascular dropout related to density of drusen in human eyes with early age-related macular degeneration. Invest. Ophthalmol. Vis. Sci. 52, 1606–1612. 10.1167/iovs.10-6476

Mullins, R.F., Schoo, D.P., Sohn, E.H., Flamme-Wiese, M.J., Workamelahu, G., Johnston, R.M., Wang, K., Tucker, B.A., Stone, E.M., 2014. The membrane attack complex in aging human choriocapillaris: relationship to macular degeneration and choroidal thinning. Am. J. Pathol. 184, 3142–3153. 10.1016/j.ajpath.2014.07.017

Pons, M., Marin-Castaño, M.E., 2011. Nicotine increases the VEGF/PEDF ratio in retinal pigment epithelium: a possible mechanism for CNV in passive smokers with AMD. Invest. Ophthalmol. Vis. Sci. 52, 3842–3853. 10.1167/iovs.10-6254

Saigal, K., Salama, J.E., Pardo, A.A., Lopez, S.E., Gregori, N.Z., 2025. Modifiable Lifestyle Risk Factors and Strategies for Slowing the Progression of Age-Related Macular Degeneration. Vis. Basel Switz. 9, 16. 10.3390/vision9010016

Sancho, M., Leiva, D., Lucendo, E., Orzáez, M., 2022. Understanding MCL1: from cellular function and regulation to pharmacological inhibition. FEBS J. 289, 6209–6234. 10.1111/febs.16136

Schlecht, A., Leimbeck, S.V., Jägle, H., Feuchtinger, A., Tamm, E.R., Braunger, B.M., 2017. Deletion of Endothelial Transforming Growth Factor-β Signaling Leads to Choroidal Neovascularization. Am. J. Pathol. 187, 2570–2589. 10.1016/j.ajpath.2017.06.018

Seo, E., Kang, H., Choi, H., Choi, W., Jun, H.-S., 2019. Reactive oxygen species-induced changes in glucose and lipid metabolism contribute to the accumulation of cholesterol in the liver during aging. Aging Cell 18, e12895. 10.1111/acel.12895

Sharma, A., Patil, J.A., Gramajo, A.L., Seigel, G.M., Kuppermann, B.D., Kenney, C.M., 2012. Effects of hydroquinone on retinal and vascular cells in vitro. Indian J. Ophthalmol. 60, 189–193. 10.4103/0301-4738.95869

Sharma, A., Yuen, D., Huet, O., Pickering, R., Stefanovic, N., Bernatchez, P., de Haan, J.B., 2016. Lack of glutathione peroxidase-1 facilitates a pro-inflammatory and activated vascular endothelium. Vascul. Pharmacol. 79, 32–42. 10.1016/j.vph.2015.11.001

Sharma, R., George, A., Nimmagadda, M., Ortolan, D., Karla, B.-S., Qureshy, Z., Bose, D., Dejene, R., Liang, G., Wan, Q., Chang, J., Jha, B.S., Memon, O., Miyagishima, K.J., Rising, A., Lal, M., Hanson, E., King, R., Campos, M.M., Ferrer, M., Amaral, J., McGaughey, D., Bharti, K., 2021. Epithelial phenotype restoring drugs suppress macular degeneration phenotypes in an iPSC model. Nat. Commun. 12, 7293. 10.1038/s41467-021-27488-x

Sherman, B.T., Hao, M., Qiu, J., Jiao, X., Baseler, M.W., Lane, H.C., Imamichi, T., Chang, W., 2022. DAVID: a web server for functional enrichment analysis and functional annotation of gene lists (2021 update). Nucleic Acids Res. 50, W216–W221. 10.1093/nar/gkac194

Silverstein, R.L., Febbraio, M., 2009. CD36, a scavenger receptor involved in immunity, metabolism, angiogenesis, and behavior. Sci. Signal. 2, re3. 10.1126/scisignal.272re3

Skerka, C., Lauer, N., Weinberger, A.A.W.A., Keilhauer, C.N., Sühnel, J., Smith, R., Schlötzer-Schrehardt, U., Fritsche, L., Heinen, S., Hartmann, A., Weber, B.H.F., Zipfel, P.F., 2007. Defective complement control of factor H (Y402H) and FHL-1 in age-related macular degeneration. Mol. Immunol. 44, 3398–3406. 10.1016/j.molimm.2007.02.012

Sohn, E.H., Flamme-Wiese, M.J., Whitmore, S.S., Workalemahu, G., Marneros, A.G., Boese, E.A., Kwon, Y.H., Wang, K., Abramoff, M.D., Tucker, B.A., Stone, E.M., Mullins, R.F., 2019. Choriocapillaris Degeneration in Geographic Atrophy. Am. J. Pathol. 189, 1473–1480. 10.1016/j.ajpath.2019.04.005

Somasundaran, S., Constable, I.J., Mellough, C.B., Carvalho, L.S., 2020. Retinal pigment epithelium and age-related macular degeneration: A review of major disease mechanisms. Clin. Experiment. Ophthalmol. 48, 1043–1056. 10.1111/ceo.13834

Song, M.J., Quinn, R., Nguyen, E., Hampton, C., Sharma, R., Park, T.S., Koster, C., Voss, T., Tristan, C., Weber, C., Singh, A., Dejene, R., Bose, D., Chen, Y.-C., Derr, P., Derr, K., Michael, S., Barone, F., Chen, G., Boehm, M., Maminishkis, A., Singec, I., Ferrer, M., Bharti, K., 2023. Bioprinted 3D outer retina barrier uncovers RPE-dependent choroidal phenotype in advanced macular degeneration. Nat. Methods 20, 149–161. 10.1038/s41592-022-01701-1

Suchting, S., Freitas, C., le Noble, F., Benedito, R., Bréant, C., Duarte, A., Eichmann, A., 2007. The Notch ligand Delta-like 4 negatively regulates endothelial tip cell formation and vessel branching. Proc. Natl. Acad. Sci. U. S. A. 104, 3225–3230. 10.1073/pnas.0611177104

Sweigard, J.H., Yanai, R., Gaissert, P., Saint-Geniez, M., Kataoka, K., Thanos, A., Stahl, G.L., Lambris, J.D., Connor, K.M., 2014. The alternative complement pathway regulates pathological angiogenesis in the retina. FASEB J. Off. Publ. Fed. Am. Soc. Exp. Biol. 28, 3171–3182. 10.1096/fj.14-251041

Tang, S., Yang, J., Xiao, B., Wang, Y., Lei, Y., Lai, D., Qiu, Q., 2024. Aberrant Lipid Metabolism and Complement Activation in Age-Related Macular Degeneration. Invest. Ophthalmol. Vis. Sci. 65, 20. 10.1167/iovs.65.12.20

ten Brink, S.C.A., Koolen, L., Klaver, C.C.W., Bakker, R.A., den Hollander, A.I., Almedawar, S., 2025. Non-canonical roles of CFH in retinal pigment epithelial cells revealed by dysfunctional rare CFH variants. Stem Cell Rep. 20, 102385. 10.1016/j.stemcr.2024.11.015

Tzoumas, N., Hallam, D., Harris, C.L., Lako, M., Kavanagh, D., Steel, D.H.W., 2021. Revisiting the role of factor H in age-related macular degeneration: Insights from complement-mediated renal disease and rare genetic variants. Surv. Ophthalmol. 66, 378–401. 10.1016/j.survophthal.2020.10.008

Velilla, S., García-Medina, J.J., García-Layana, A., Dolz-Marco, R., Pons-Vázquez, S., Pinazo-Durán, M.D., Gómez-Ulla, F., Arévalo, J.F., Díaz-Llopis, M., Gallego-Pinazo, R., 2013. Smoking and age-related macular degeneration: review and update. J. Ophthalmol. 2013, 895147. 10.1155/2013/895147

Voigt, A.P., Mulfaul, K., Mullin, N.K., Flamme-Wiese, M.J., Giacalone, J.C., Stone, E.M., Tucker, B.A., Scheetz, T.E., Mullins, R.F., 2019. Single-cell transcriptomics of the human retinal pigment epithelium and choroid in health and macular degeneration. Proc. Natl. Acad. Sci. U. S. A. 116, 24100–24107. 10.1073/pnas.1914143116

Voigt, A.P., Mullin, N.K., Stone, E.M., Tucker, B.A., Scheetz, T.E., Mullins, R.F., 2021. Single-cell RNA sequencing in vision research: Insights into human retinal health and disease. Prog. Retin. Eye Res. 83, 100934. 10.1016/j.preteyeres.2020.100934

Wang, H., Yang, Y., Ye, Y., Wei, X., Chen, S., Cheng, B., Lv, Y., 2025. Srxn1 Overexpression Protect Against Cardiac Remodelling by Inhibiting Oxidative Stress and Inflammation. J. Cell. Mol. Med. 29, e70432. 10.1111/jcmm.70432

Wang, L., Clark, M.E., Crossman, D.K., Kojima, K., Messinger, J.D., Mobley, J.A., Curcio, C.A., 2010. Abundant lipid and protein components of drusen. PloS One 5, e10329. 10.1371/journal.pone.0010329

Wang, Y., Fung, N.S.K., Lam, W.-C., Lo, A.C.Y., 2022. mTOR Signalling Pathway: A Potential Therapeutic Target for Ocular Neurodegenerative Diseases. Antioxidants 11, 1304. 10.3390/antiox11071304

Wang, Y., Ma, H., Ge, J., Chen, Kuangqi, Chen, Kai, Ye, H., Pan, X., Wang, X., Xia, J., Shen, J., Cheng, T., Cui, H., Sheng, Y., 2026. Senescence-induced endothelial-to-mesenchymal transition accelerates the subretinal fibrosis in neovascualr age-related macular degeneration. J. Transl. Med. 24, 251. 10.1186/s12967-026-07707-z

Whitmore, S.S., Braun, T.A., Skeie, J.M., Haas, C.M., Sohn, E.H., Stone, E.M., Scheetz, T.E., Mullins, R.F., 2013. Altered gene expression in dry age-related macular degeneration suggests early loss of choroidal endothelial cells. Mol. Vis. 19, 2274–2297.

Xu, X., Pang, Y., Fan, X., 2025. Mitochondria in oxidative stress, inflammation and aging: from mechanisms to therapeutic advances. Signal Transduct. Target. Ther. 10, 190. 10.1038/s41392-025-02253-4

Yang, Y., Dieter, M.Z., Chen, Y., Shertzer, H.G., Nebert, D.W., Dalton, T.P., 2002. Initial characterization of the glutamate-cysteine ligase modifier subunit Gclm(-/-) knockout mouse. Novel model system for a severely compromised oxidative stress response. J. Biol. Chem. 277, 49446–49452. 10.1074/jbc.M209372200

Yoon, H., Shaw, J.L., Haigis, M.C., Greka, A., 2021. Lipid metabolism in sickness and in health: Emerging regulators of lipotoxicity. Mol. Cell 81, 3708–3730. 10.1016/j.molcel.2021.08.027

Yue, X., Liang, S., Zhang, H., Dai, T., Wu, J., 2025. Association of Decay Accelerating Factor (CD55) Positive Extracellular Vesicles with Advanced Age and Blood Glucose Levels in Elderly Individuals. Clin. Interv. Aging 20, 1549–1560. 10.2147/CIA.S536479

Zauhar, R., Biber, J., Jabri, Y., Kim, M., Hu, J., Kaplan, L., Pfaller, A.M., Schäfer, N., Enzmann, V., Schlötzer-Schrehardt, U., Straub, T., Hauck, S.M., Gamlin, P.D., McFerrin, M.B., Messinger, J., Strang, C.E., Curcio, C.A., Dana, N., Pauly, D., Grosche, A., Li, M., Stambolian, D., 2022. As in Real Estate, Location Matters: Cellular Expression of Complement Varies Between Macular and Peripheral Regions of the Retina and Supporting Tissues. Front. Immunol. 13, 895519. 10.3389/fimmu.2022.895519

Zuo, J., Pan, Y., Wang, Y., Wang, W., Zhang, H., Zhang, S., Wu, Y., Chen, J., Yao, Q., 2025. ROS-responsive drug delivery system with enhanced anti-angiogenic and anti-inflammatory properties for neovascular age-related macular degeneration therapy. Mater. Today Bio 32, 101757. 10.1016/j.mtbio.2025.101757

