## Supplementary material for "Oxidative Stress Susceptibility, Complement Dysregulation, and Metabolic Reprogramming in *CFH* Y402H Patient-Derived Choriocapillaris Endothelial Cells"

Majlinda Lako

Newcastle University

Biosciences Institute

International Centre for Life

Central Parkway

Newcastle NE1 3BZ

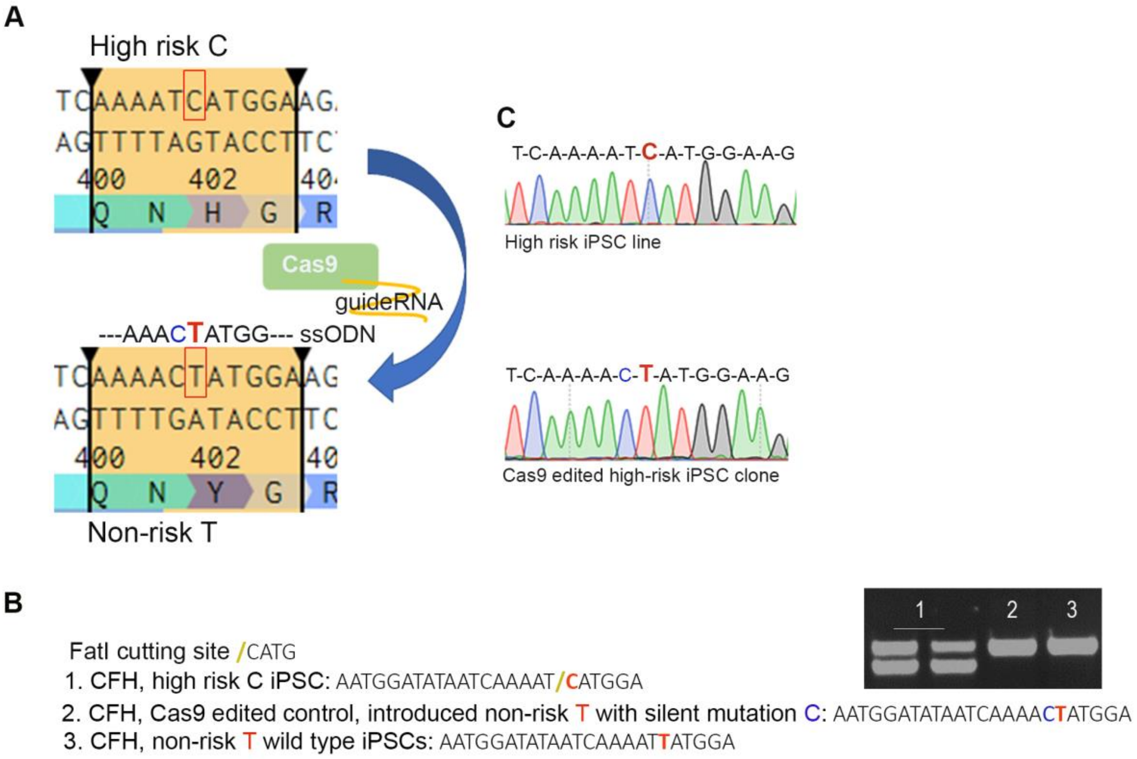


**Figure S1. Correction of high-risk (HR) C nucleotide to T (H402Y) in the *CFH* gene. A**) Schematic representation of CRISPR/Cas9 editing of high-risk iPSCs, substituting C to T (in red rectangle). **B**) Clone screening method using *FatI* restriction enzyme cutting at 'CATG depicted in yellow. C nucleotide changed to T, depicted in red, a silent mutation in blue. **C**) Example of DNA sequencing of HR iPSCs and the CRISPR/Cas9 edited isogenic control clone with non-risk T and silent mutation.

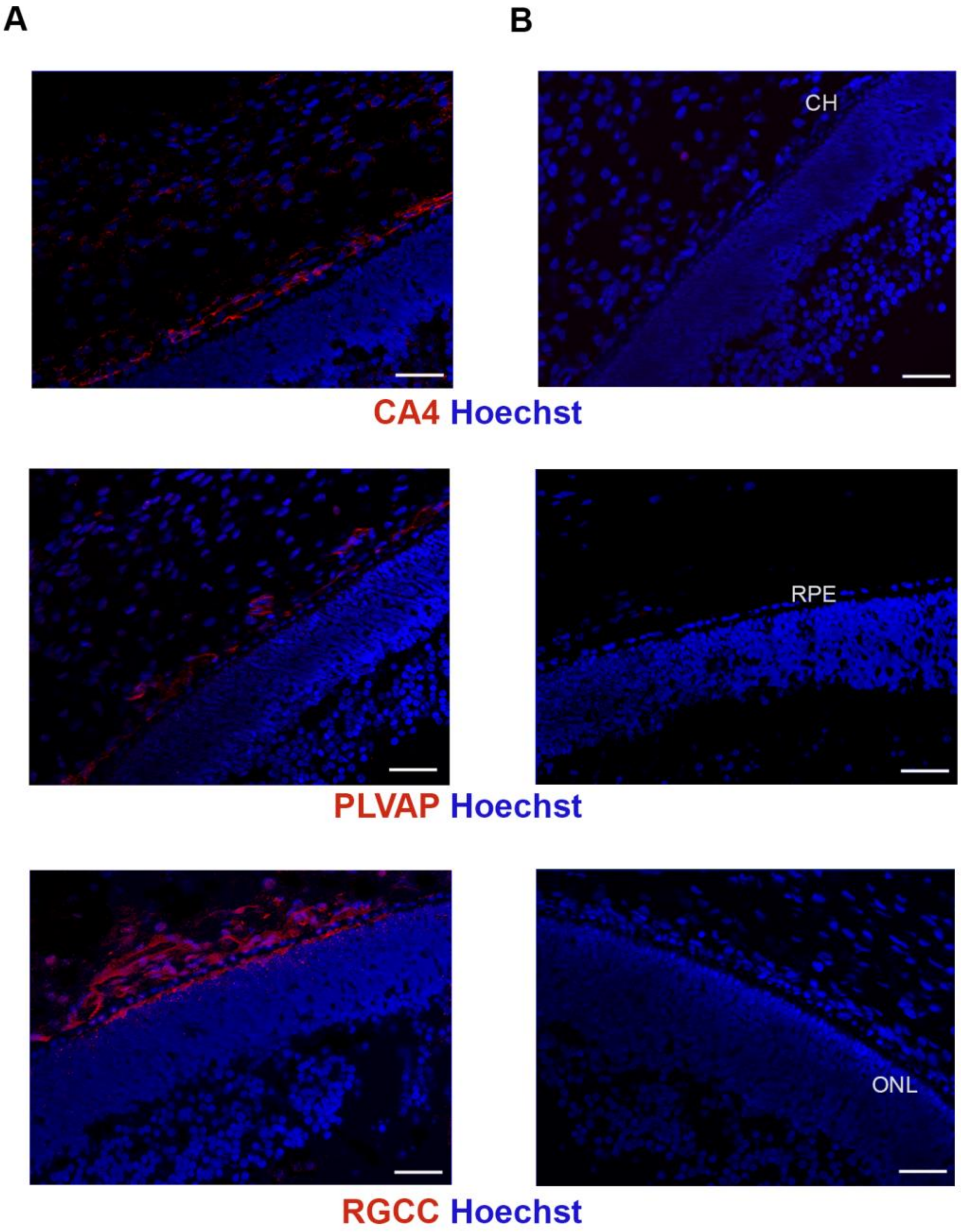


**Figure S2. Optimisation of CEC-specific markers immunocytochemical staining using 12 PCW eye sections**. CA4, PLVAP, and RGCC are depicted in red (**A**), nuclei were counterstained with Hoechst, secondary antibody-only control is shown in (**B**).

Scale bars = 50 µm. CH: choroid, RPE: retinal pigment epithelium, ONL: outer nuclear layer.

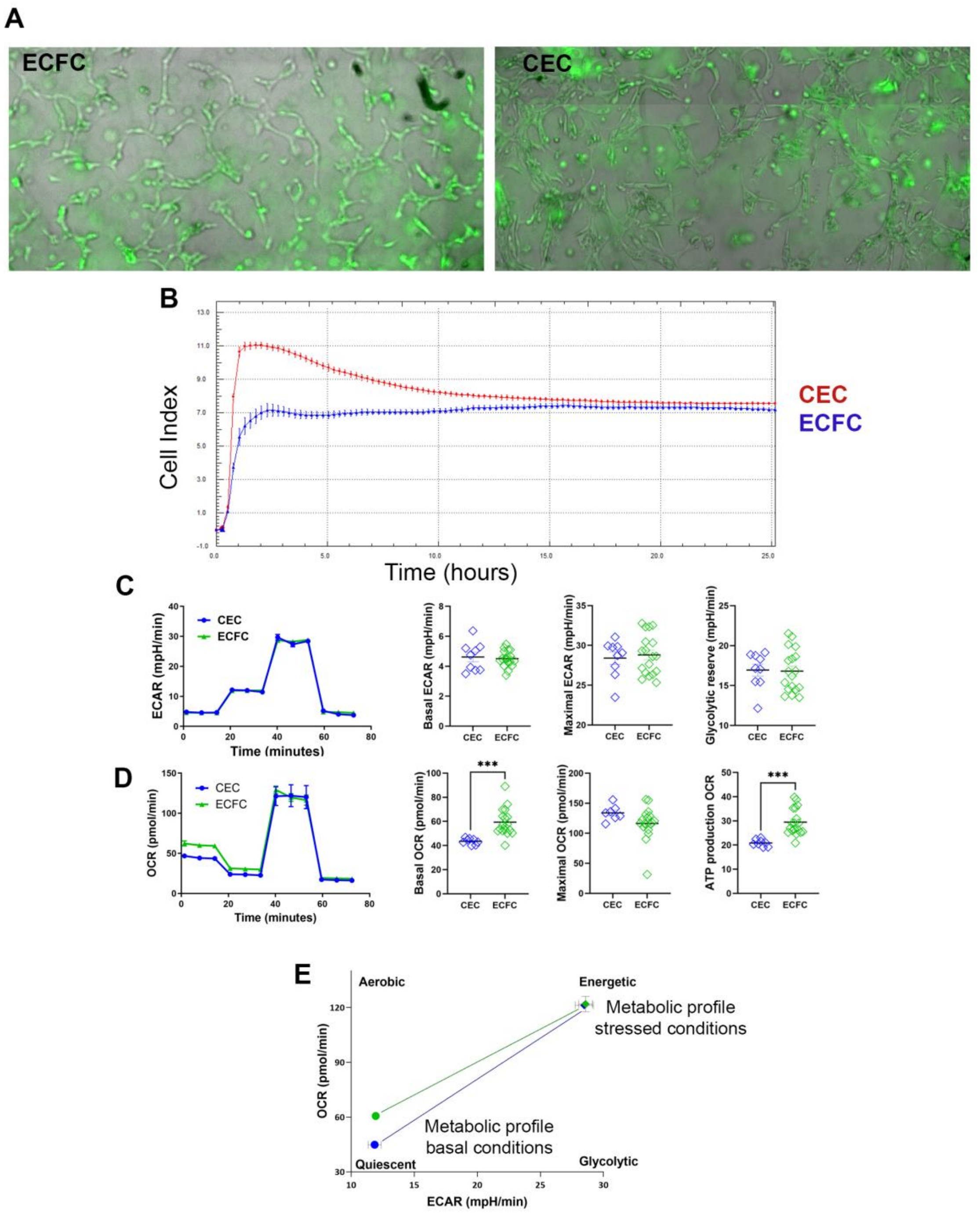


**Figure S3**. **Functional and metabolic analyses demonstrate no significant differences between iPSC-CECs and ECFCs**. **A**) Tubulogenesis assay for iPSC-derived CECs compared to ECFCs; Calcein images were merged with brightfield. **B**) Barrier capacity, measured as cell index; a higher cell index indicates higher barrier capacity. **C**) Seahorse glycolysis: This panel depicts cell glycolytic activity as measured by the Seahorse analyser, which assesses the extracellular acidification rate (ECAR). **D**) Mitochondrial respiration: This panel shows measurements of cellular mitochondrial activity via the oxygen consumption rate (OCR). **E**) Energy map: This panel combines ECAR and OCR data to present a "cellular energy phenotype," categorising the metabolic state of the cells. The map typically plots ECAR (glycolytic activity) versus OCR (mitochondrial respiration), showing relative contributions and shifts between energy pathways under different conditions (e.g., basal and stressed states). Data shown as mean ± SEM, n = 8, unpaired t-test.

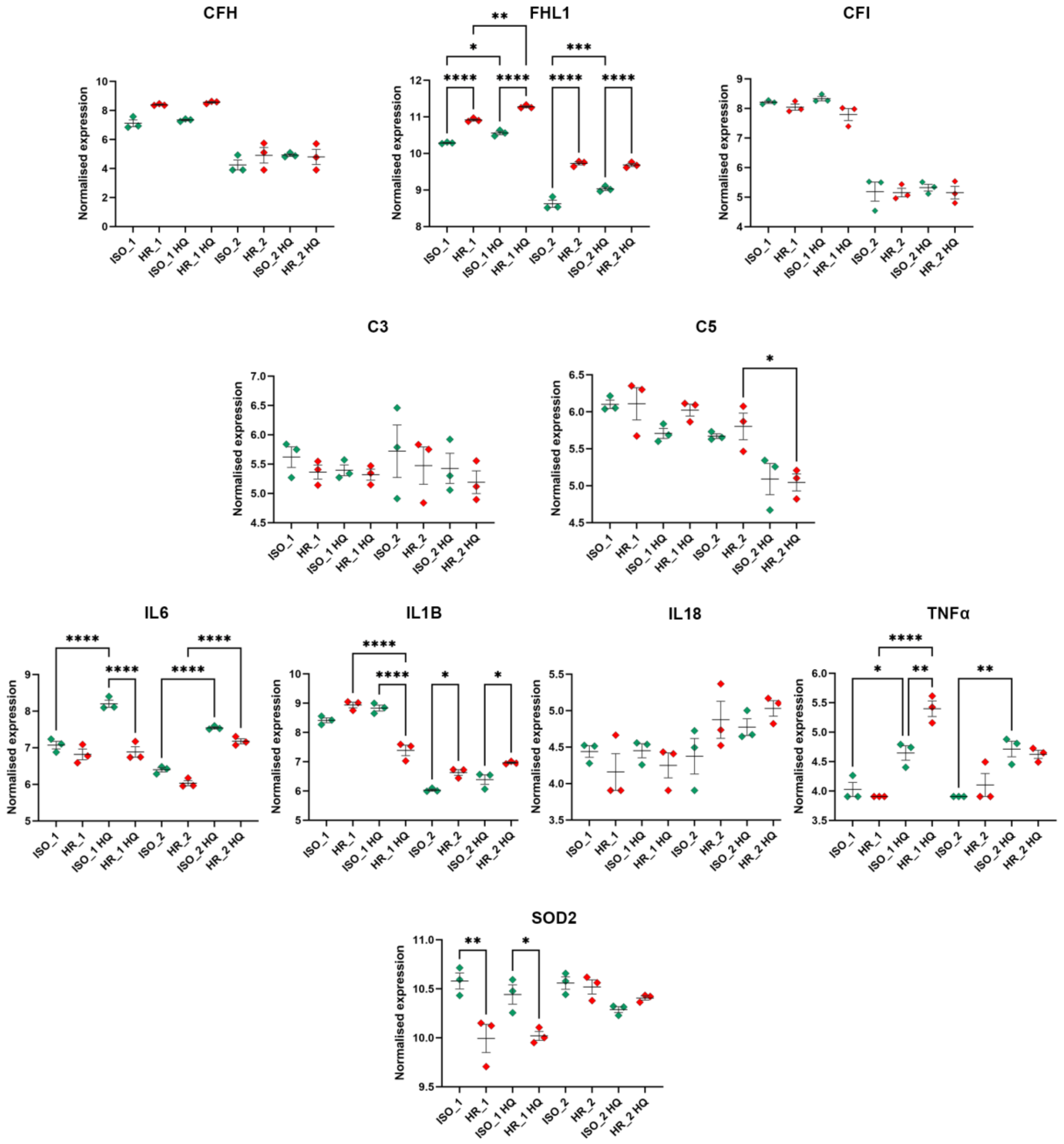


**Figure S4. *CFH, FHL1, CFI, C3, C5, IL6, IL1B, IL18, TNFα,* and *SOD2* mRNA normalised expression from the bulk RNA sequencing dataset (DESeq2)**. Data are presented as mean ± SEM (n=3), one-way ANOVA, * $p < 0.05$, ** $p < 0.01$, *** $p < 0.001$, and **** $p < 0.0001$.

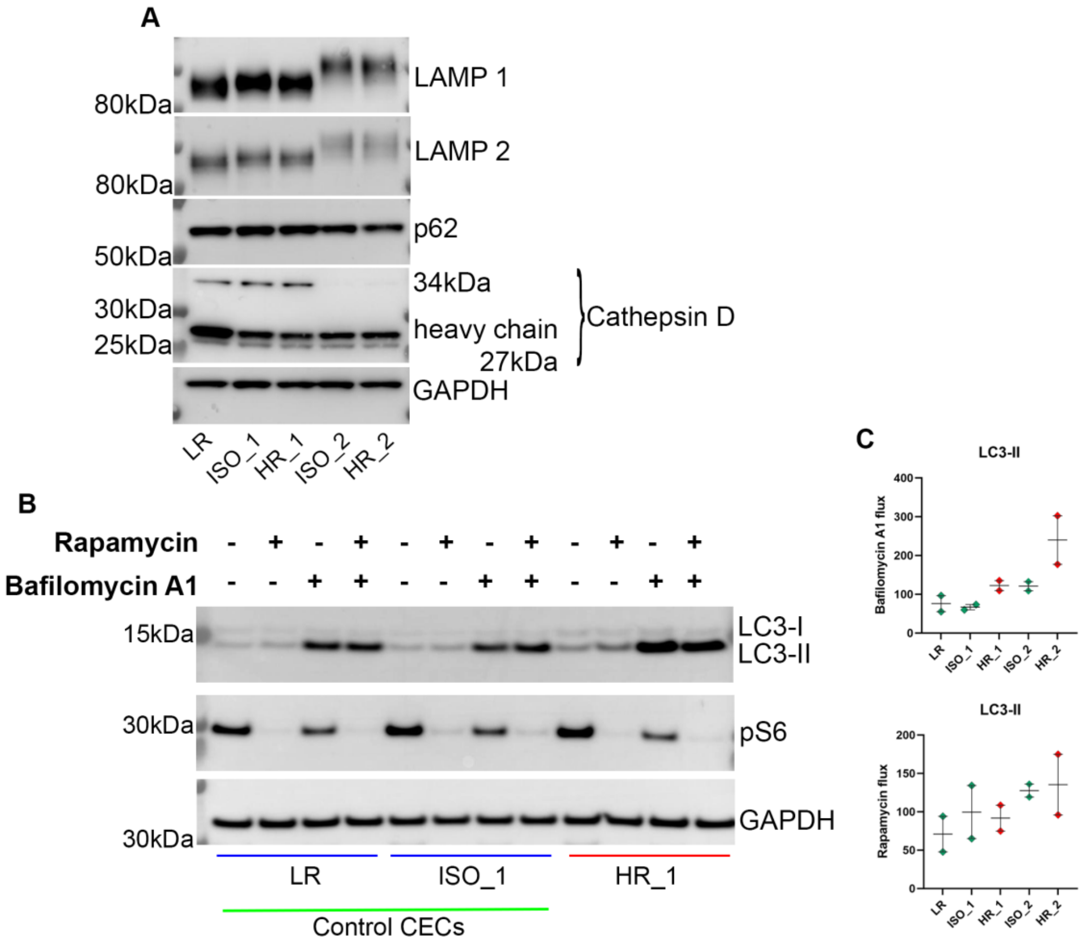


**Figure S5. Autophagic flux in HR- versus ISO- and LR-CECs**. **A**) Representative western blot steady-state levels of autophagy markers. **B**) Representative western blot images depicting LC3 and p62 in CECs treated with Bafilomycin A1, preventing formation of autolysosomes (24 hours) and/or Rapamycin, inducing autophagosome formation. GAPDH was used as a loading control. **C**) Graphs present both rapamycin and bafilomycin LC3-II flux rates. Rapamycin flux was calculated by subtracting the LC3-II band value for double treatment from the single rapamycin treatment. Bafilomycin flux was generated by subtracting the single bafilomycin treatment from the basal untreated control. Data were normalised to GAPDH expression and presented as mean ± SEM (n=2). No significant differences were detected between HR- and ISO-CECs by two-way ANOVA. LR: low-risk CECs.

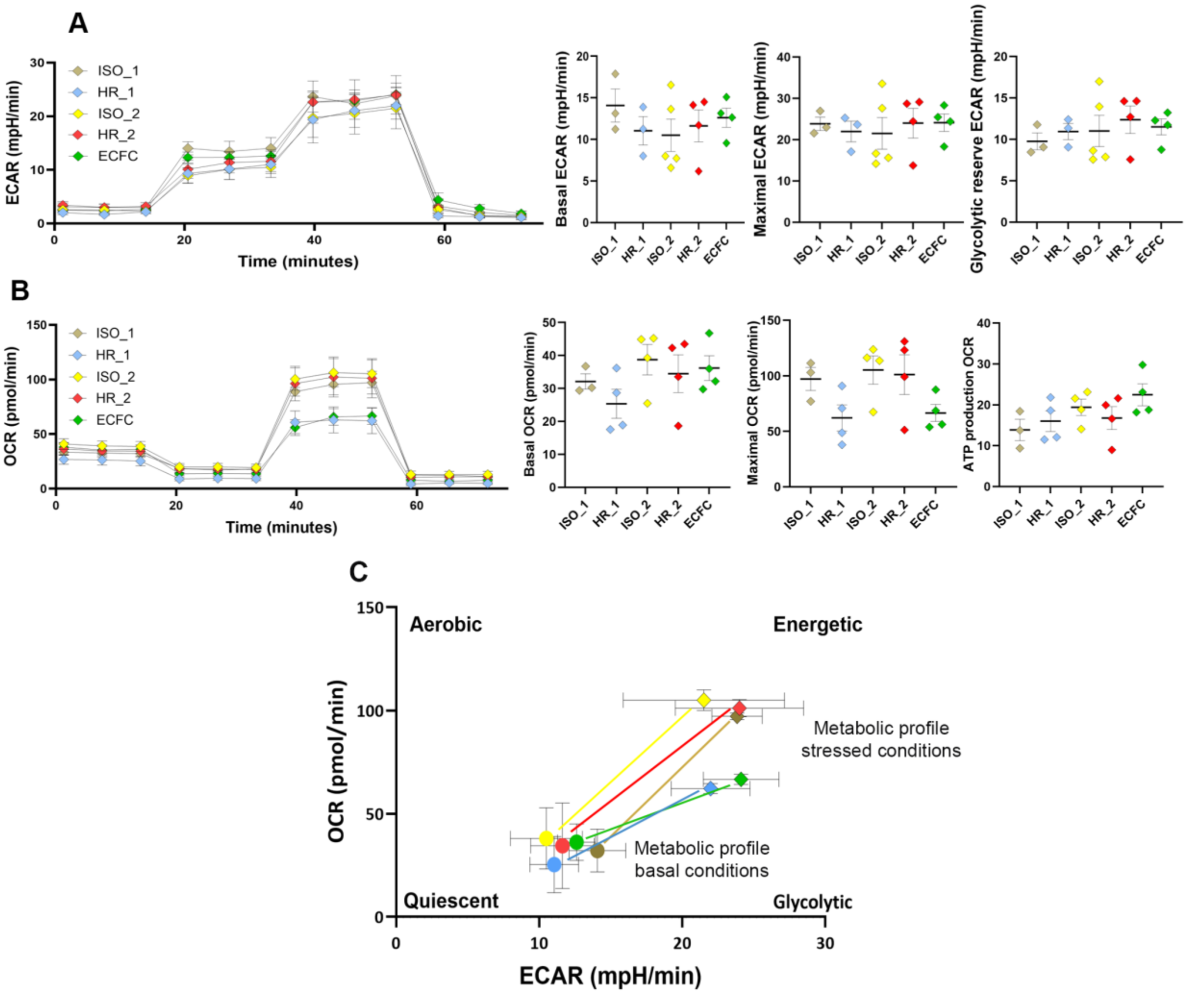


**Figure S6**. **High-risk and isogenic control CECs glycolysis (A), mitochondrial respiration levels (B) and Energy map (C).** No significant differences were observed between HR- and ISO-CECs. Data shown as mean ± SEM, n = 3, one-way ANOVA.

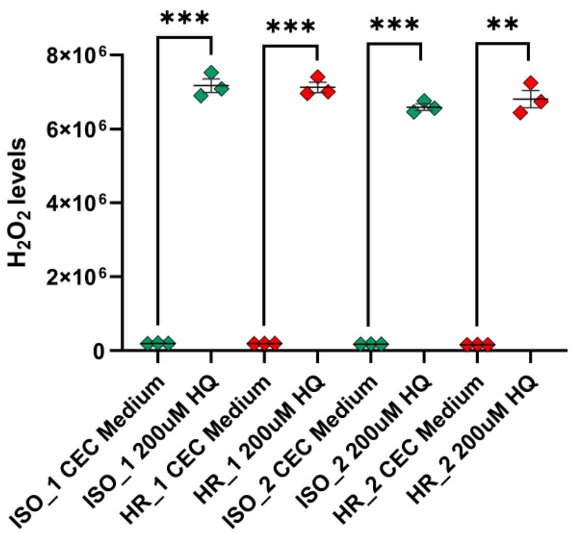


**Figure S7. $H_2O_2$ levels in HR and ISO CECs in the presence of 200 µM HQ in comparison to control conditions**. Data are presented as mean ± SEM (n=3), unpaired t-test, ** p < 0.01, *** p < 0.001.

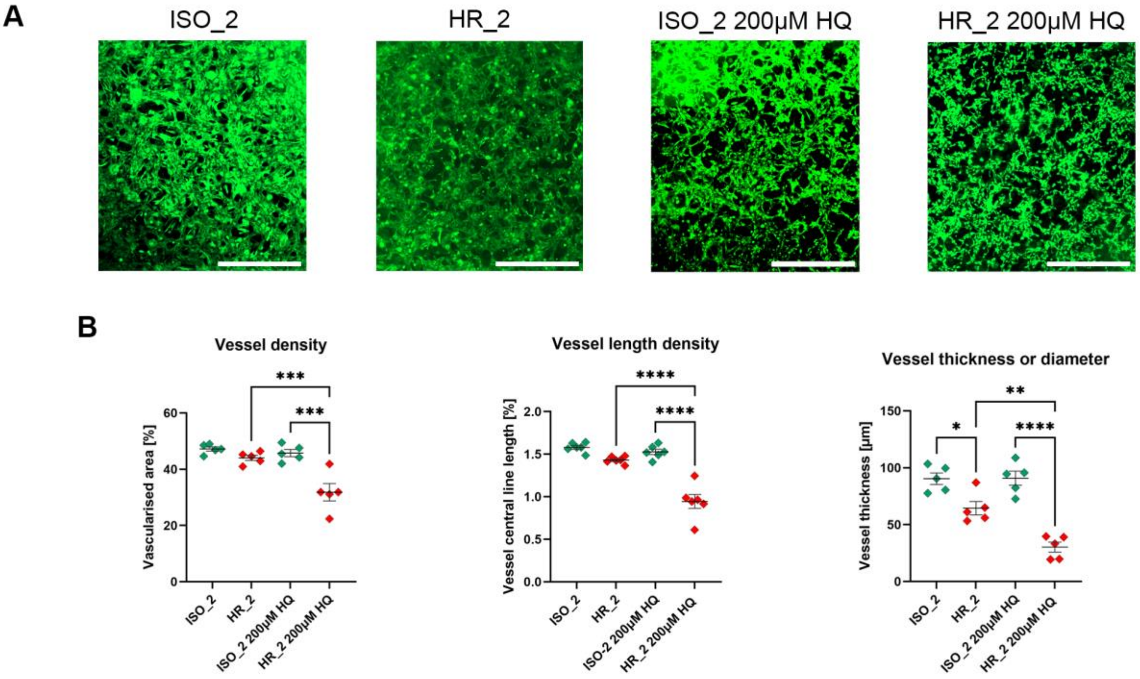


**Figure S8**. **HR_2 and ISO_2 endothelial tube formation ability in the presence of 200 µM hydroquinone**. **A**) Representative images of HR and its ISO. Scale Bar: 1000 µm. **B**) The graphs show quantified vessel density, vessel length and vessel thickness for control and HQ conditions. Data shown as mean ± SEM, n = 5, one-way ANOVA, * p < 0.05, ** p < 0.01, *** p < 0.001, and **** p < 0.0001.

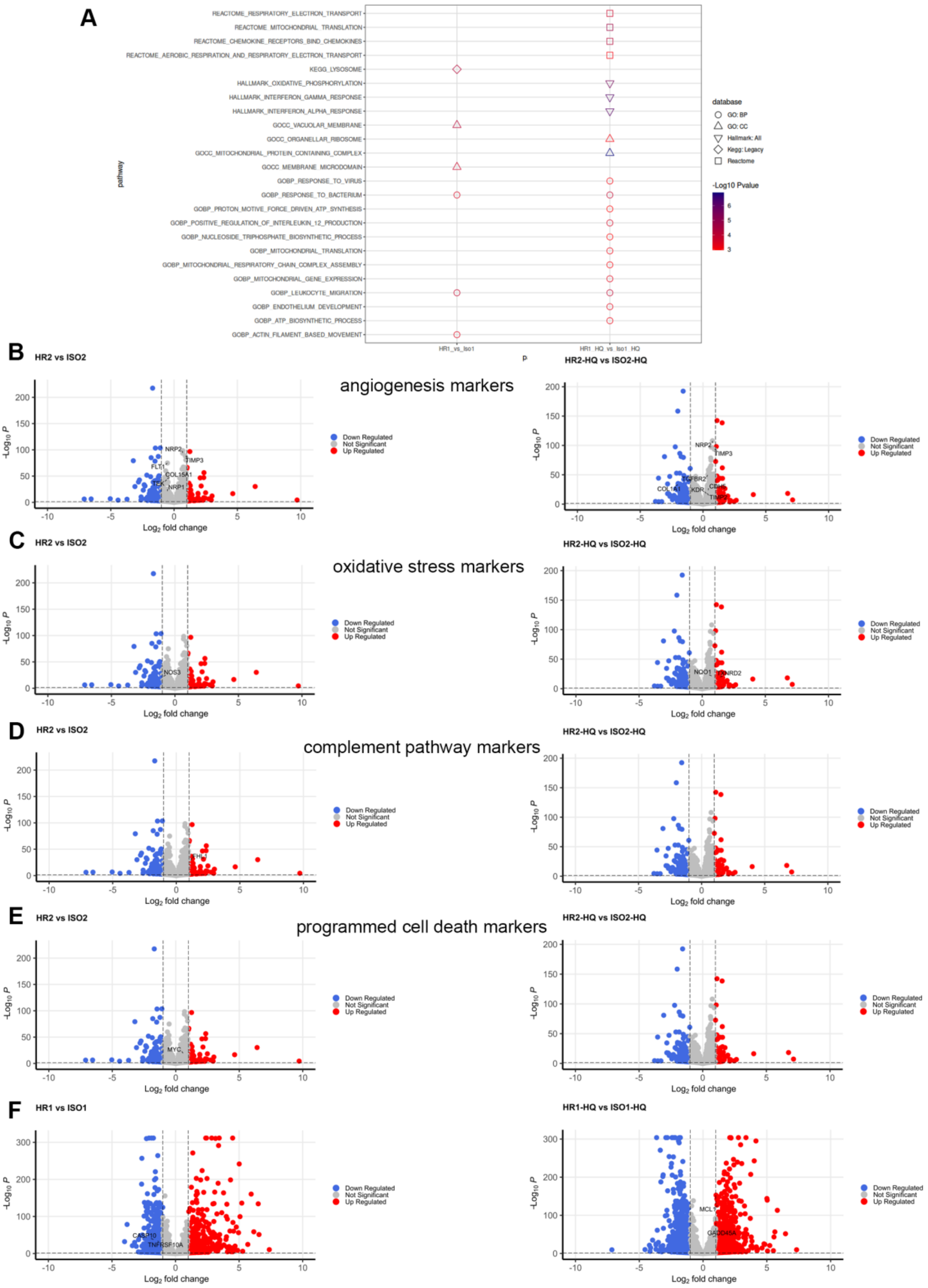


**Figure S9**. **A**) GO, Hallmark, KEGG and Reactome analyses of downregulated DEGs in HR_1 in comparison to ISO_1 CECs with or without the HQ. Volcano plots depicting angiogenesis (**B**), oxidative stress (**C**), complement pathway (**D**) and programmed cell death

(**E**) marker gene expression in HR_2 cells versus ISO_2 under HQ and control conditions. **F**) Programmed cell death marker gene expression in HR_1 cells under HQ and control conditions. DEG: differentially expressed gene. DEGs were identified from relevant contrasts using DESeq2. Significance thresholds shown, absolute $Log_2$ fold change of 1 and adjusted p value < 0.05.

| Primers | Off-target sites for gRNA: AATGGATATAATCAAAATCA **TGG** correcting high-risk C nucleotide to T (H402Y) in the *CFH* gene to create isogenic controls | Sequencing result for CRIPR/Cas9 generated isogenic controls |
|---|---|---|
| **F: CTGACTCGTGTCCCAGGTG**<br>**R: TGGAATAGTGTCTTCTCCCATACA** | **I.** AATGcAaAgAATCAAAATCA **TGG (chr2, + strand)** | AAAATGCAAAGAATCAAAATCA**TGG**TT |
| **F: GCATACGTACATACCAGGCCA**<br>**R: TTTAGGCTGCAGTTTCTCAAAAT** | II. AATGGATATAAagtAAATCA **TGG (chr5, + strand)** | AAAATGGATATAAAGTAAATCA**TGG**CC |
| **F: GCTTTAAGACAGCCATGAAAGTGA**<br>**R: ACAGTGTTATCACTCCCCCA** | III. AAaGGcTAaAATCAAAATCA **CGG (chr7, + strand)** | AAAAAGGCTAAAATCAAAATCA**CGG**AT |

**Table S1. Off-target sequences for H402Y in the *CFH* gene.**

| Target | Dilution | Supplier | Catalogue number |
| --- | --- | --- | --- |
| CA4 | ICC: 1:100 | R&D Systems | MAB21861 |
| CASP3 | ICC: 1:400 | Cell Signaling Technology | 9661S |
| CD31 | ICC: 1: 300 | DAKO | M0823 |
| C3 | WB: 1:500 | Abcam | ab48611 |
| activated C3 (C3b, iC3b, and C3c) | ICC: 1:50 | Hycult Biotech | HM2168 |
| C5b-9 | ICC: 1:200 | Agilent | M0777 |
| CFH | WB: 1:1000 | Merck | 341276 |
| CFI | WB: 1:250 | LSBio | LS-C147802 |
| CTSD | WB: 1:2000<br>ICC: 1:200 | Sigma-Aldrich | C0715 |
| GAPDH | WB: 1:5000 | Santa Cruz Biotechnology | sc-47724 |
| LAMP1 | WB: 1:500 | Developmental Studies Hybridoma Bank | H4A3 |
| LAMP2 | WB: 1:500<br>ICC: 1:100 | Abcam | ab199946 |
| LC3 | WB: 1:500 | Cell Signaling Technology | 3868S |
| p62 | WB: 1:500<br>ICC: 1:200 | BD Transduction Lab | 610832 |
| p-S6 (Ser235/236) | WB: 1:500 | Cell Signaling Technology | 2211 |
| PLVAP | ICC: 1:100 | Sigma-Aldrich | HPA002279 |
| RGCC | ICC: 1:50 | abcam | AB221098 |

**Table S2. List of primary antibodies used for western-blot and immunofluorescence analysis.**

| Gene name | Primers | AT |
| --- | --- | --- |
| CD31 | F: CCAAGGTGGGATCGTGAGG<br>R: TCGGAAGGATAAAACGCGGTC | 60 |
| RGCC | F: CACTGTCACTCCTCAGAAAGCTAA<br>R: TGTCCCCTCTGGCAGCAGAT | 60 |
| PLVAP | F: GCTGCTGGTATTACCTGCG<br>R: GCCATAGACCATGAAGAGCAC | 60 |

**Table S3. Primers for quantitative RT-PCR.**

**Table S4**. Differential gene expression between HR_1/2 and ISO_1 /2 in the presence of 200 μM HQ or control conditions. DEGs were identified from relevant contrasts using DESeq2, with a gene characterised as differently expressed if the contrast gave an absolute $Log_2$ fold change of 1 and an adjusted p-value < 0.05.

**Table S5**. GO/Hallmark/KEGG/Reactome enrichment of DEGs listed for HR_1/2 in comparison to ISO_1/2 in the presence of 200 μM HQ or control conditions. Gene Set Enrichment Analyses (GSEA) were completed using the R packages clusterProfiler and fgsea, where possible, the top 20 contrasts were selected for visualisation based on ascending p-value.
